# Pre-existing genome architecture partitions viral takeover in the smallest free-living eukaryote

**DOI:** 10.64898/2026.10.29.762491

**Authors:** Keith Harrison, Martha Valiadi, Estelle S. Kilias, Yann Loe-Mie, Ramy Ragheb, Susan Kimmance, Adam Monier

## Abstract

Nuclear DNA viruses colonize host genomes already structured into complex spatial domains, yet whether established spatial boundaries dictate viral progression remains unresolved—in part because sequence composition and transcriptional output are intertwined in animal models. Here, we tracked productive lytic infection of the marine picoeukaryote *Ostreococcus tauri* by the giant double-stranded DNA prasinovirus OtV5, recording transcription, accessibility, and 3D contact dynamics across parallel time courses. Although total cellular transcript pools collapsed 18-fold by 10 h post-infection, the host transcriptional shutdown, decay of insulation, early chromatin opening, and viral DNA positioning were all bounded by architectural coordinates defined prior to viral arrival. Across two low-GC outlier domains, chromatin accessibility spiked rapidly, spatial insulation eroded, and loci exhibited relative upregulation by 6 hpi, with the smaller outlier region retaining elevated transcript levels per cell compared to the rest of the genome. Unexpectedly, viral DNA was preferentially depleted from these accessible outlier cores, contacting instead the standard-GC telomeric flanks situated immediately beyond the pre-existing spatial boundaries. Domain identity, rather than local accessibility or active host expression, thus governs viral positioning, demonstrating that pre-existing nuclear architecture partitions the physical and transcriptional takeover of the host cell.

## Introduction

Nuclear DNA viruses take over host genomes already folded into complex three-dimensional domains that differ in primary sequence, epigenetic state, transcriptional output, and nuclear localization (1). Although viral entry can remodel these features while viral DNA interacts non-randomly with host chromatin (2), global population assays typically average across these genomic subdivisions. Consequently, whether viral takeover progresses uniformly or is guided by architectural boundaries established before infection remains unresolved. Disentangling this relationship is key to linking host transcriptional shutdown with physical nuclear reorganization.

The marine picoeukaryote *Ostreococcus tauri* (class Mamiellophyceae, order Mamiellales), the smallest free-living eukaryote at ∼0.8 μm (3), offers an ideal model to address this question. Its compact nuclear genome spans 20 chromosomes, including the big outlier chromosome (BOC, chromosome 2) and small outlier chromosome (SOC, chromosome 19). Both harbor low-GC, transposon-dense, hypomethylated outlier regions (BOC1 and SOC1) enriched for stress-response loci, metabolic functions, and structural rearrangements (4–7). Chromosome conformation capture in uninfected cells established that BOC1 and SOC1 form spatially insulated domains bounded by defined coordinates (7), and a gapless telomere-to-telomere assembly resolves these outlier regions completely (8). Analogous outlier domains are conserved in the other Mamiellales genus *Bathycoccus* (9). Lytic large double-stranded DNA prasinoviruses (phylum *Nucleocytoviricota*, family Phycodnaviridae) such as OtV5 routinely infect natural *Ostreococcus* populations (10–12). Lacking an encoded RNA polymerase (11), OtV5 depends on nuclear machinery during infection (13), placing host and viral genomes within the same nuclear space where transcription, physical interactions, and chromatin accessibility can be tracked in parallel.

OtV5 infection induces widespread host transcriptional shutdown alongside upregulation of host histone variants and methyltransferases, suggesting chromatinisation of viral DNA inside the host nucleus (13), though physical contacts and genome architecture were not examined. Other members of Phycodnaviridae actively alter host chromatin structure: for example, the chlorovirus PBCV-1 packages the histone H3 lysine 27 methyltransferase vSET to suppress host gene expression (14). In animal viruses, actively transcribed genomes target accessible host chromatin (15, 16), whereas latent Epstein-Barr virus episomes and inactive hepatitis B virus covalently closed circular DNA localize to repressive nuclear compartments (17–20). However, composition and transcriptional activity are confounded in animal models because repressive compartments are simultaneously AT-rich and silenced (2). While infection can restructure host 3D architecture (21–25) and Hi-C has mapped integrated proviruses within host boundaries (26, 27), whether pre-existing domain architecture structures the host response to productive lytic infection in a microbial eukaryote is unknown. BOC1 and SOC1 enable a direct test because their distinctive sequence composition and spatial insulation exist prior to viral infection.

Here we asked how viral takeover alters host expression and spatial genome architecture, and whether pre-existing domains constrain this reorganisation. We performed paired RNA-seq, Hi-C, and ATAC-seq across infection time courses to measure host and viral transcription, chromatin accessibility, local insulation, and inter-genome contact dynamics simultaneously. Host transcription collapsed 18-fold per cell, yet the response was spatially partitioned along pre-existing insulation boundaries that bounded the transcriptional change, insulation decay, early chromatin opening, and viral DNA positioning. OtV5 infection thus decouples compartment identity from transcriptional activity and accessibility, two factors that remain confounded in animal systems.

## Results

### OtV5 collapses host transcription across the genome

We exposed *O. tauri* RCC4221 populations to OtV5 during the initial light phase and demonstrated viral entry by electron microscopy (Fig. 1A). Progression matched documented lytic dynamics (13): cellular disruption emerged between 6 and 8 hpi, generating a burst size of 13.6 ± 1.9 virions per cell. Host population clearance reached 97.3 ± 0.26% by 24 hpi, concurrent with a 2.7-fold elevation in extracellular viral particles above pre-infection levels (Fig. 1B; data S1).

**Fig. 1.**
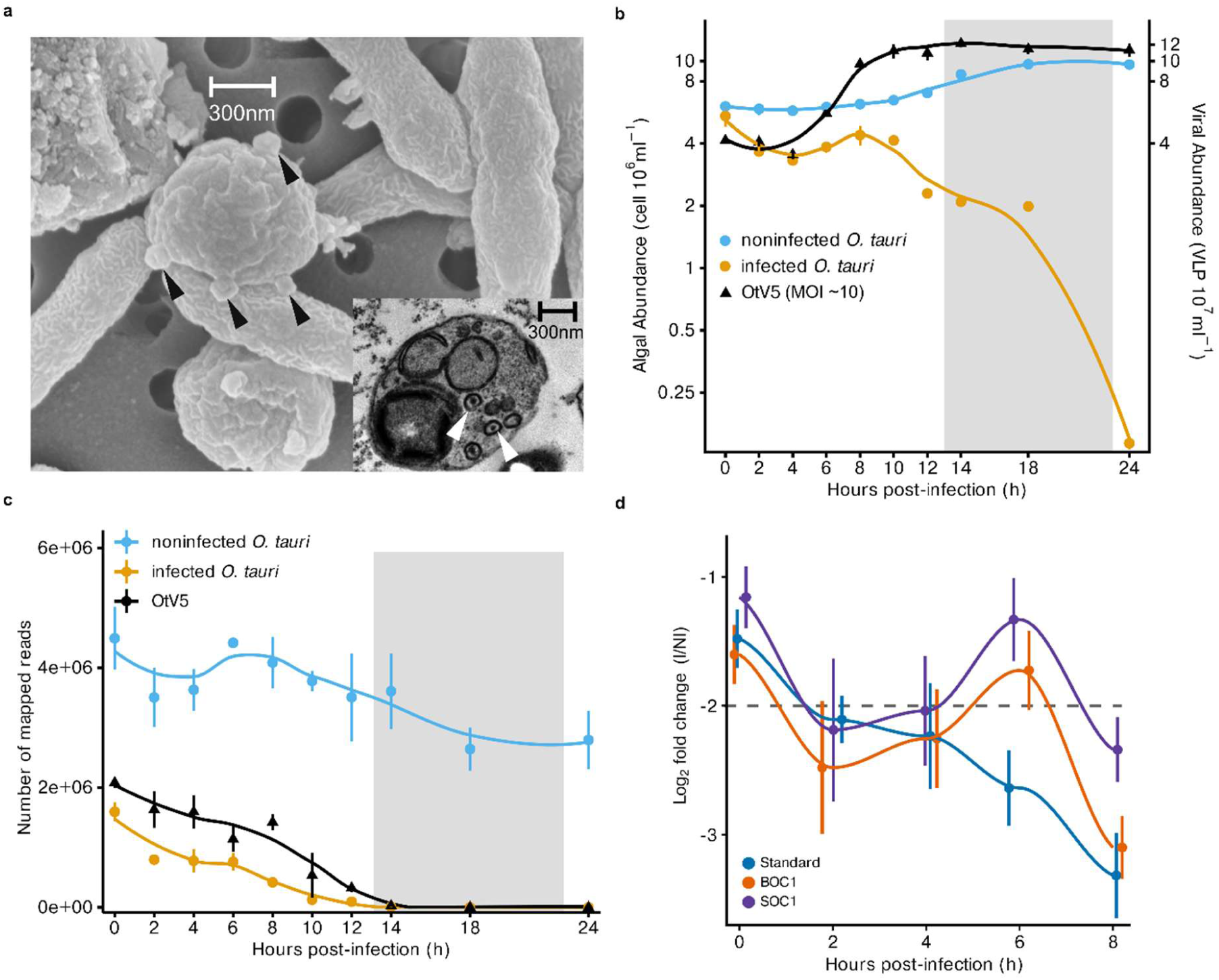
OtV5 infection dynamics and the genome-wide host transcriptional collapse. (**A**) Representative scanning electron micrograph of an OtV5-infected *O. tauri* culture with a transmission electron micrograph inset. Large spheres are algal cells, small spheres are virions (arrowheads), and rod-shaped structures are bacteria. Scale bars, 300 nm. (**B**) Flow-cytometric quantification of algal abundance (cells mL-1, left axis) and viral abundance (virus-like particles mL-1, right axis) in infected and noninfected cultures across 24 hpi (OtV5; MOI = 9 by flow cytometry). The shaded box (13 to 23 h) marks the dark period. Data are means ± SD (*n* = 4 biological replicates). (**C**) Total mapped host and viral transcript reads across the infection time course (means ± SD, *n* = 4 biological replicates). Per-cell normalised values are given in fig. S1. (**D**) Log2 fold change in *O. tauri* transcript abundance (infected versus noninfected) over time, with genes grouped by encoding region: Standard, BOC1 (chromosome 2 outlier region), and SOC1 (chromosome 19 outlier region). Data are means ± SD (*n* = 4 biological replicates); the dashed line marks log2FC = -2. Non-outlier regions of BOC and SOC are shown in fig. S2B.

Population breakdown capped a global transcriptional shutdown triggered at inoculation. Uninfected cultures contained 4,472,999 ± 522,983 host transcript reads at 0 hpi, whereas infected cultures displayed reduced host mRNA pools (1,591,433 ± 160,721) alongside abundant viral transcription (2,080,101 ± 88,643). Host output reached near-complete cessation by 10 hpi (130,768 ± 94,260 reads), contrasting with sustained transcript levels in uninfected controls (3,777,592 ± 170,599; Fig. 1C).

Raw sequence counts fail to distinguish host transcriptional decay from cell loss or sequencing space competition; standard compositional normalisation calculates relative fractions rather than absolute cellular yield, mistaking absolute repression for relative induction during global collapse (28–32). Standardising host reads against cell density via hourly flow cytometry confirmed the transcript loss (fig. S1). Per-cell host transcript pools dropped 2.4-fold at 0 hpi and 18.5-fold by 10 hpi relative to controls (log2FC -1.29 to - 4.21; Welch’s *t* test, *P*adj < 0.01; fig. S1). Cell density adjustments account for part of this decrease, reducing log2 fold shifts by 0.20 to 1.62 units. Viral RNA displacement fails to explain the remaining loss: under fixed pool sizes, expected host reads at 0 hpi equal 1,938,829, whereas observed reads fell 17.9% lower (1,591,433). Host transcription is thus repressed at infection onset beyond sequencing or cell count artifacts. Because viral transcription had already initiated, the infected 0 hpi sample represents an early stage of viral takeover rather than an uninfected baseline, leaving uninfected controls as the sole reference state.

Transcriptional shutdown proceeded non-uniformly across the nuclear genome. Outlier regions maintained higher mean transcript levels than standard genomic regions throughout infection (Welch’s *t* test, *P*adj < 0.01; Fig. 1D; fig. S2; tables S1 and S2), prompting an examination of the boundaries governing this transcriptional partition.

Viral transcription exhibited a regulated temporal cascade typical of Phycodnaviridae (33, 34). OtV5 genes resolved into three kinetic classes based on peak expression: early (0 hpi, *n* = 103), mid (2 hpi, *n* = 31), and late (4 to 8 hpi, *n* = 112) (Fig. 2, A and B; data S2). Early loci encoded initiation factors, topoisomerases, ligases, DNA polymerase III, SF3 helicases, and RNA processing factors, indicating rapid co-optation of host transcription machinery rather than chromosomal cleavage (Fig. 2C). Mid-phase loci governed nucleotide synthesis and replication machinery, dominated by OtV5_154c, a putative UBOX-domain ubiquitin ligase factor (Fig. 2C; data S2). Late loci encoded structural proteins, major capsids, and glycosyltransferases. Promoter motif analysis identified distinct upstream elements for early (CCTAAGT, 25 sites, E-value = 2.8e-5) and late classes (bipartite AAAAATCT-----TATA-TA, 64 sites, E-value = 2.1e-38; Fig. 2D), whereas mid-phase elements lacked strong support (CTTTAA, 14 sites, E-value = 7.3e-1). OtV5 expression is thus transcriptionally orchestrated, imposing viral expression kinetics upon a collapsing host transcriptome.

**Fig. 2.**
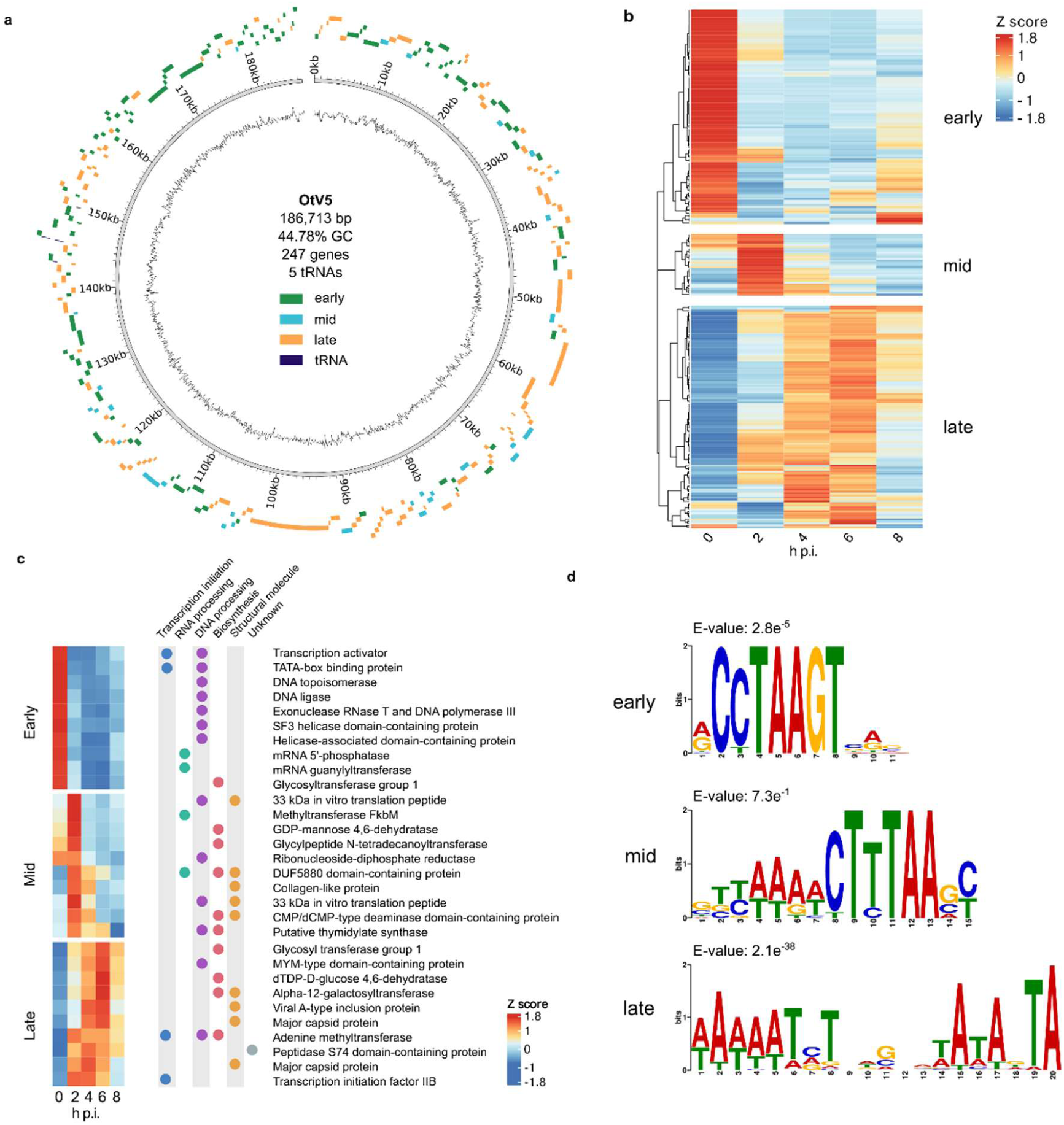
Viral gene expression kinetics and regulatory architecture of OtV5. (**A**) Circular map of the OtV5 genome with genes coloured by kinetic class (early, mid, late) and the inner track showing GC content. (**B**) Expression heatmap of all viral genes across infection timepoints (0, 2, 4, 6, 8 hpi), normalised per gene (z score) and hierarchically clustered. (**C**) Mean expression of the ten most highly expressed annotated genes in each kinetic class, centred and scaled per timepoint; functional categories are indicated by the coloured circles. (**D**) Upstream promoter motifs for each kinetic class, identified by differential enrichment (MEME); sequence logos and E-values are shown (early, 25 sites; mid, 14 sites; late, 64 sites).

Infection disrupted normal host diel expression rhythms. Non-metric multidimensional scaling (NMDS) of control transcriptomes displayed the expected cyclic path, whereas infected samples diverged from this cycle by 2 hpi (Fig. 3A). Non-negative matrix factorisation (NMF) of uninfected control data resolved four transcriptional modules matching G1, S, G2, and M phases (fig. S3, A and B; data S3), each tied to specific physiological functions (data S4). Standard chromosomes showed high G2 activity (>40% per chromosome), whereas outlier regions followed alternative schedules: BOC1 expression spanned G2, M, and G1 phases (37%, 24%, and 29%), and SOC1 exhibited primary G1 alignment (49%) (fig. S3C; data S3). The outlier domains operated on a distinct transcriptional schedule prior to viral contact, providing the baseline against which infection-driven reorganization was measured.

**Fig. 3.**
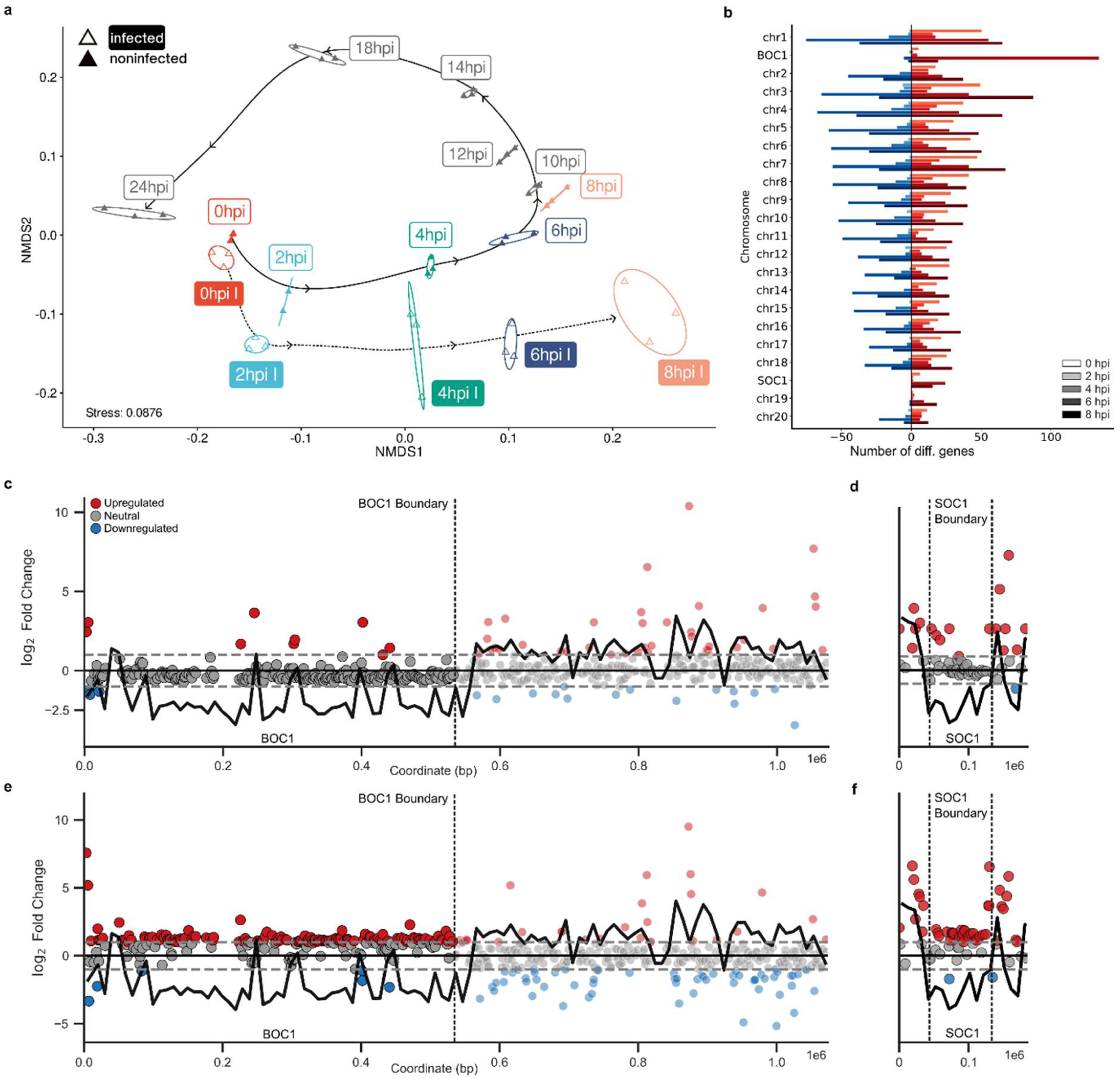
The host transcriptional response terminates at a pre-existing architectural boundary. (**A**) Non-metric multidimensional scaling of host transcriptomes across all timepoints and both infection states; samples are coloured by infection status (filled symbols, noninfected; open symbols, infected) with outline and shape colour denoting timepoint. Stress value and sample size are shown on the plot. (**B**) Number of differentially expressed genes (|log2FC| > 1, adjusted *P* < 0.1) per chromosome across timepoints (0 to 8 hpi, indicated by colour depth); genes called up and down are shown in red and blue. BOC1 and SOC1 are plotted as rows separate from their parent chromosomes. Calls are made relative to library composition and are interpreted as relative rather than absolute change (see Results). (**C** to **F**) Gene-level log2 fold change (infected versus noninfected) along chromosomal coordinates for BOC (C, E) and SOC (D, F) at 2 hpi (C, D) and 6 hpi (E, F). Dotted lines mark the outlier-region boundaries mapped in noninfected cells (7); the bold black line shows %GC in 10 kb windows. The relative transcriptional response at 6 hpi is confined within these boundaries.

### Pre-existing genomic compartments partition the collapse of host transcription

To dissect this non-uniform transcriptional decay at gene-level resolution, we evaluated infected against uninfected host transcriptomes across time, profiling differential output chromosome by chromosome (Fig. 3B). Infection onset triggered a muted host response, with outlier domains appearing largely unperturbed: only 12 out of 591 differentially expressed loci (|log2FC| > 1, adjusted *P* < 0.1) mapped within BOC1 or SOC1 (data S5). Initial upregulated transcripts encoded nucleic acid-binding factors, DNA helicases, DNA polymerase B, condensin subunits, and histone variants H2B and H3 (data S5 and S6; figs. S4 and S5). Furthermore, a cohort of signal-peptide-bearing factors, including an elicitor-inducible LRR receptor-like protein and a superoxide dismutase, was distributed across standard chromosomes rather than localized to outlier regions (table S3). Notably, two immediate transcriptional shifts occurred on outlier chromosomes: the BOC1-localized H2A.Z histone variant (OT_ostta02g00100) experienced fast repression, whereas an SNF2/INO80-family chromatin remodeling factor displayed increased expression (fig. S4; data S6).

By 6 hpi, this transcriptional landscape had inverted completely. Standard chromosomes exhibited wide-scale bidirectional regulation (599 upregulated and 905 downregulated loci; data S7), whereas BOC1 and SOC1 featured exclusively upregulated transcripts without a single downregulated call (*n* = 134 on BOC1, *n* = 33 on SOC1; Fig. 3B; data S7).

Because DESeq2 (35) normalizes against total library size, these statistical calls reflect changes relative to a rapidly collapsing global baseline. When interpreted alongside per-cell measurements, loci classified as upregulated on BOC1 at 6 hpi represent transcripts whose cellular abundance decays significantly slower than the host mean, rather than genes experiencing true de novo transcriptional induction. We refer to this behavior as relative upregulation throughout. These relatively upregulated loci were strictly compartmentalized in space. On BOC, relative upregulation extended continuously from the chromosomal terminus to a precise junction at ∼535 kb, beyond which expression dynamics matched standard genomic regions (Fig. 3, C and E). Crucially, this coordinate was not inferred post hoc from infected datasets; it mirrors the exact insulation boundary identified in uninfected *O. tauri* (7). Thus, pre-existing spatial architecture delineates the physical boundary of the host transcriptional response. This feature is also recovered in a recent gapless telomere-to-telomere RCC4221 genome (8), which completely resolves duplicated sequences within outlier loci, recovers the identical breakpoint. SOC1 exhibited matching domain-restricted dynamics, with relative upregulation restricted to its core outlier region rather than its standard-GC telomeric flanks (Fig. 3, D and F).

Integrating regional expression proportions (tables S1 and S2) with absolute per-cell transcript counts (fig. S1) clarifies the functional impact of this structural partition. Every nuclear region experiences absolute transcript depletion during takeover; no domain preserves baseline expression. Through 4 hpi, output across all three genomic sectors declined at comparable rates (log2 fold shifts -1.1 to -1.7), after which their trajectories diverged. By 10 hpi, transcript pools from standard chromosomes and BOC1 dropped 18-fold (log2 -4.2) and 17-fold (log2 -4.1), respectively, whereas SOC1 suffered only a 9-fold reduction (log2 -3.2), maintaining approximately double the per-cell transcript yield of standard loci (fig. S1). Consequently, SOC1 progressively expanded its relative contribution to total host output over the infection course (mean increase +5.66 percentage points; Wilcoxon signed-rank *P* = 0.002), while the output fractions of BOC1 and standard regions shrank (-4.23 and -3.37 percentage points; P = 0.014 and P = 0.037; shares are tracked over five region classes and therefore do not sum to zero). The sharpness of this boundary varies between chromosomes. On BOC, the standard-GC arm (nonBOC1) mirrors standard genomic decay almost perfectly, separating a resistant domain from a typical chromosomal arm. On chromosome 19, however, the standard-GC flank (nonSOC1) exhibits decay dynamics closer to SOC1 than to standard chromosomes (fig. S1), indicating that transcriptional retention spills over beyond the SOC1 core. Because nonSOC1 contains few annotations, this requires cautious interpretation; nevertheless, the functional boundary is sharp on BOC and transitional on SOC. Overall, pre-existing spatial architecture defines a sharp shift in expression behavior while absolute transcript abundance declines globally.

Genes exhibiting relative retention encompassed stress mediators, repair machinery, and central metabolic pathways. The most prominent retained locus on BOC1 at 6 hpi encoded a SUMO peptidase; additional retained factors across BOC1 and SOC1 involved cellular stress and DNA repair systems (HSP70, Clp protease, protein phosphatase 2C, SNF2/INO80, RAD21), protein synthesis and folding apparatus (eIF translation factors, methyltransferases, serine/threonine protein kinases, ER retention receptors, nascent polypeptide-associated complexes), and key metabolic enzymes (glycerol-3-phosphate O-acyltransferase, a giant type 1 polyketide synthase) (data S7). BOC1 also harbors a conserved suite of Mamiellales genes dedicated to nucleic acid metabolism, including a sister chromatid cohesion factor, Tc1-like transposases, a PHD-finger protein, and the replication licensing factor MCM3, several of which showed altered transcript levels during infection (fig. S4). Notably, RNA polymerase III subunits situated on BOC1 displayed relative upregulation, including Rpc3 (OT_ostta02g00600), RPC10 (OT_ostta02g01780), and core structural components OT_ostta02g01770 and OT_ostta02g02290 (figs. S4 and S6; data S5 and S7). This spatial partition also shaped post-transcriptional regulation: alternative splicing events preferentially accumulated within BOC1 and SOC1 throughout infection, remaining largely static across the rest of the host genome (fig. S7; data S8).

### The same boundary partitions chromatin architecture

Should this expression boundary reflect underlying 3D organization, structural remodeling must be detectable during active viral takeover. We mapped chromosome conformation via Hi-C in control and infected cells at 2 and 6 hpi (10 kb resolution; data S9; fig. S8). Distance-dependent decay rates remained virtually identical between states (fig. S9), confirming that global nuclear folding persists during infection.

Beneath this preserved macro-architecture, outlier domains diverged markedly from standard genomic tracks. Evaluating differential insulation scores across regions (Fig. 4A) revealed that standard chromosomes maintained stable insulation throughout infection, whereas BOC1 and SOC1 experienced pronounced insulation shifts relative to standard loci (*P* < 0.05, two-sided Mann-Whitney *U* test with FDR correction; Fig. 4A). Track inspection mapped this remodeling precisely: insulation values within BOC1 and the SOC1 core increased by 2 and 6 hpi, signaling progressive decay of spatial insulation, whereas adjacent standard-GC flanks and SOC telomeres remained unaffected (Fig. 4, B and C; fig. S10). Local architectural breakdown is thus strictly localized to outlier domains, constrained by the identical coordinates that mark host transcriptional retention.

**Fig. 4.**
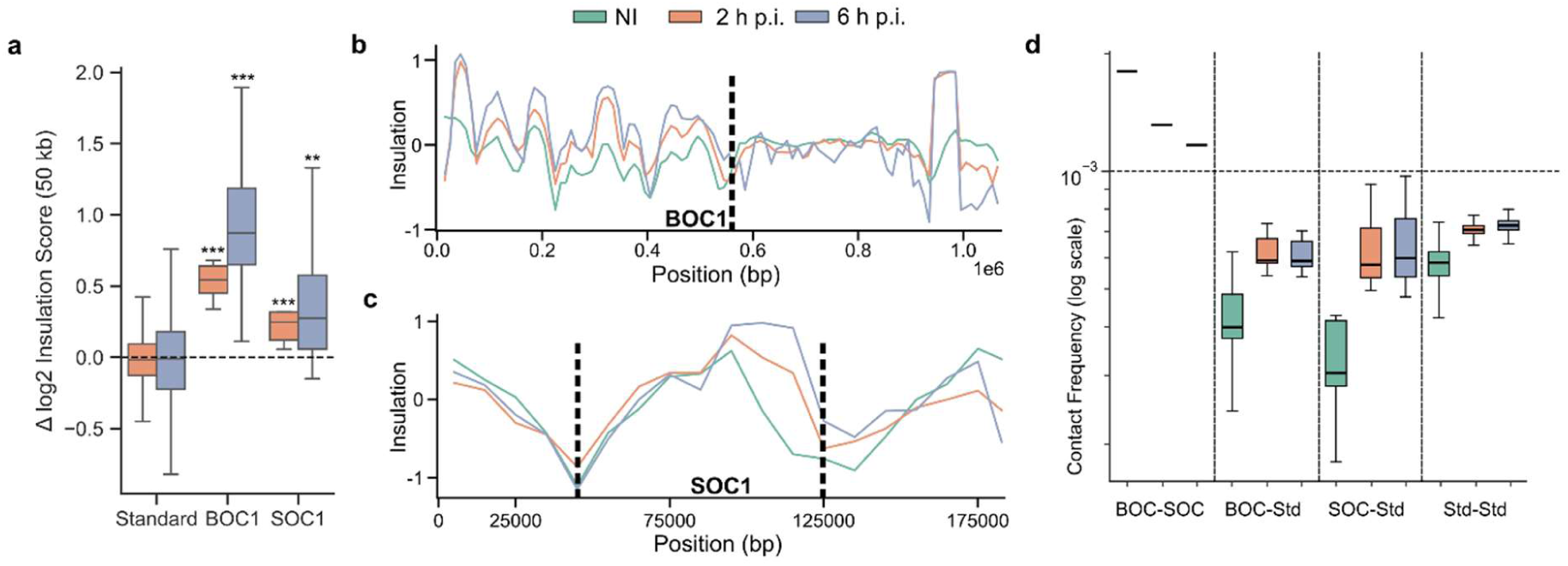
The same boundary partitions chromatin architecture during infection. (**A**) Change in log2 insulation score (Δ, 50 kb window around 10 kb bins) at 2 and 6 hpi relative to noninfected cells, for Standard regions (*n* = 1222 bins) and the outlier regions BOC1 (*n* = 52) and SOC1 (*n* = 9). (**B** and **C**) Insulation tracks (50 kb resolution) for noninfected, 2 hpi and 6 hpi across BOC (B) and SOC (C); outlier-region boundaries are marked by dashed lines. (**D**) Interchromosomal (trans) contact frequencies (log scale) grouped into four classes: BOC-SOC (1 region pair), BOC-standard (18 pairs), SOC-standard (18 pairs), and standard-standard (153 pairs). In (A) and (D) boxes are coloured by timepoint. In (A) significance versus the Standard background was assessed by two-sided Mann-Whitney *U* test with FDR correction (\**P* < 0.05, \*\**P* < 0.01, \*\*\**P* < 0.001). The BOC-SOC class contains a single region pair and is shown for comparison rather than as a formal test.

These compartments also displayed distinct relationships between physical contact and transcriptional output. Within BOC1, local intra-domain contact frequencies correlated positively with locus expression at 2 and 6 hpi (ρ = 0.143 and ρ = 0.167; *P* < 0.05; fig. S11; data S10), a coupling completely absent across SOC and standard chromosomes. Thus, structural density selectively supports transcriptional persistence on BOC1. Furthermore, the outlier domains exhibited strong spatial co-localization. Interchromosomal (trans) contact between BOC1 and SOC1 surpassed all other regional interactions across uninfected and infected states (Fig. 4D; data S11), exceeding baseline trans interactions with standard chromosomes by 3.11-fold in control cells, 1.87-fold at 2 hpi, and 1.63-fold at 6 hpi. Because physical distance determines contact probability (36), BOC1 and SOC1 occupy adjacent nuclear space prior to viral arrival.

This spatial association degrades during viral infection. Relative to control levels, trans-interaction frequency between BOC1 and SOC1 dropped to 0.73 by 2 hpi and 0.65 by 6 hpi, while trans contacts between outlier regions and standard chromosomes increased concurrently (Fig. 4D). Rather than acting as an impenetrable barrier, this pre-existing architectural compartment undergoes progressive physical erosion while remaining detectable through 6 hpi.

### Viral DNA is depleted from the outlier cores

Leveraging chromosome conformation capture to track inter-genomic contact dynamics, we delineated the spatial distribution of OtV5 relative to the pre-existing host architecture at 2 and 6 hpi (Fig. 5, A to C; data S12 and S13).

**Fig. 5.**
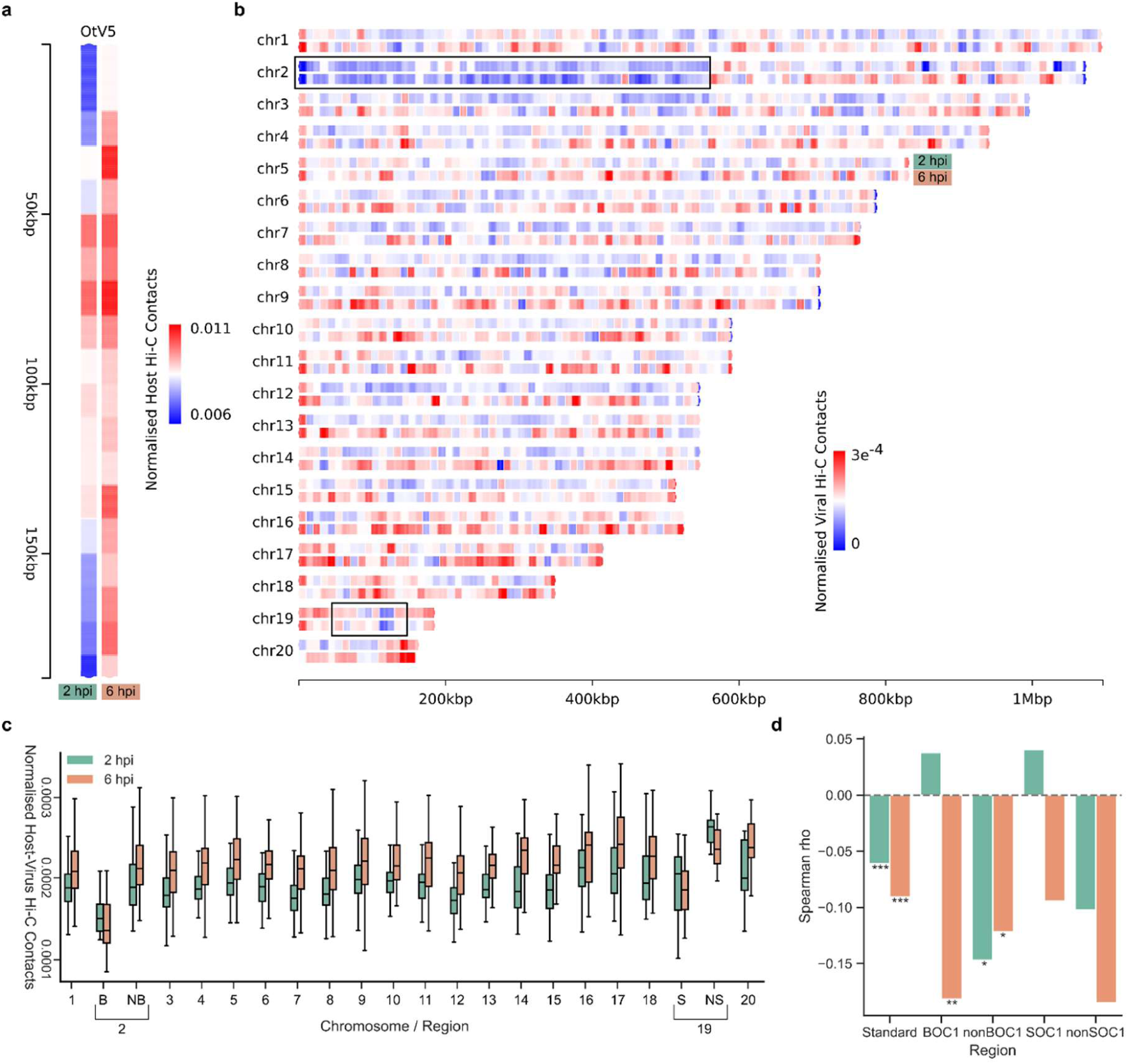
Viral DNA is depleted from the outlier cores and contacts accessible chromatin outside them. (**A**) OtV5 genome karyotype with bins coloured by their normalised Hi-C contact frequency with the entire *O. tauri* genome, at 2 and 6 hpi. (**B**) *O. tauri* genome karyotype with bins coloured by their normalised contact frequency with the OtV5 genome, at 2 and 6 hpi; outlier regions are outlined in black. (**C**) Distribution of normalised viral-host contact frequency for each host chromosome and outlier region (2 hpi, green; 6 hpi, orange). (**D**) Spearman rank-correlation coefficients (ρ) between gene expression and host-virus Hi-C contact frequency, computed across gene loci within each region at 2 hpi (green) and 6 hpi (orange). B and NB denote BOC1 and nonBOC1; S and NS denote SOC1 and nonSOC1. Asterisks denote significance of ρ (Spearman correlation test): \**P* ≤ 0.05, \*\**P* ≤ 0.01, \*\*\**P* ≤ 0.001.

Actively transcribed nuclear DNA viruses in mammalian hosts preferentially target open, highly transcribed host chromatin (15, 16). Under this paradigm, the outlier regions should attract maximum viral interaction given their sustained transcriptional output and rapid chromatin opening. Paradoxically, viral DNA contact across the host nuclear genome was spatially heterogeneous, reaching its absolute minimum within the outlier cores: BOC1 exhibited the lowest interaction frequencies of any genomic sector at both sampling points (*P* < 0.05, two-sided Mann-Whitney *U* test; Fig. 5, B and C), with exclusion intensifying as lytic takeover advanced.

Outside these depleted cores, viral DNA localization followed specific spatial preferences. At 2 hpi, interactions peaked across the accessible, GC-standard, repeat-dense SOC telomeric flanks (*P* < 0.05, Mann-Whitney *U* test; Fig. 5, B and C) situated immediately adjacent to the SOC1 architectural boundary. By 6 hpi, telomeric engagement subsided, with maximal viral contact shifting toward standard chromosomes 17 and 20 (Fig. 5, B and C; data S12 and S13). Viral positioning is thus characterized by depletion from outlier cores and temporal redistribution across standard chromatin, occupying accessible loci delineated by the identical boundary that partitions host transcription and insulation. Because these internal contrasts manifest concurrently within identical nuclei, general technical artifacts cannot account for the observed regional depletion.

The interplay between host expression and viral contact displayed a matching asymmetry. Within outlier domains, locus expression correlated inversely with viral contact density (SOC1 at 2 hpi: ρ = -0.529, *P* < 0.001; SOC1 at 6 hpi: ρ = -0.300, *P* < 0.01; BOC1 at 6 hpi: ρ = -0.182, *P* < 0.01; Fig. 5D; data S10), whereas standard chromosomes exhibited no significant relationship (ρ = -0.06 at 2 hpi; ρ = -0.09 at 6 hpi). Reciprocally, host contact profiles varied non-randomly along the viral genome (fig. S12; data S14). Evaluated in 5 kb windows, inter-genomic contacts peaked at ∼60 kb at 2 hpi and ∼57 kb at 6 hpi, mapping directly inside the FhaB gene locus (OtV5_070/OtV5_071; 54,743 to 62,876 bp). This region resides within the hypervariable genomic window diversified across prasinovirus isolates, which encodes surface-anchored FhaB proteins under positive selection and dictates host range (37). Width-matched permutation testing against a viral null distribution (10,000 iterations; fig. S12) confirmed enrichment of the hypervariable sub-window (45 to 65 kb, within the 25 to 80 kb region defined across isolates (37)) at 2 hpi (empirical *P* = 0.029) and marginal significance at 6 hpi (*P* = 0.055), whereas the enclosed FhaB locus achieved statistical significance at 6 hpi (*P* = 0.028; 2 hpi *P* = 0.33). Host interactions thus initiate broadly across the hypervariable region before refining onto FhaB loci during infection progression. Following FDR adjustment across tests (adjusted *P* = 0.058 to 0.33), we consider this spatial alignment suggestive, supported by the reproducible contact peak over an independently annotated locus across timepoints. Intriguingly, because FhaB genes belong to the late kinetic class and remain silent during early infection, host targeting occurs independently of viral transcription. A secondary contact cluster spanning 80 to 90 kb contains a PCNA-like gene adjacent to an AT-rich motif, highlighting a candidate replication origin.

### *Chromatin opens early in the outlier regions, but accessibility does not explain* low viral contact

Physical occlusion of host chromatin represents a straightforward mechanism to explain diminished viral interaction across outlier domains. To test this hypothesis and profile early nuclear remodeling, we performed ATAC-seq across noninfected diel phases (G1, S, G2, M) and hourly post-infection stages (0 to 5 hpi). Viral entry triggered a systemic loss of openness across the host genome (126 loci lost accessibility, 1 gained; |log2FC| > 0.3, adjusted *P* < 0.05; fig. S13; data S15), displaying no single-chromosome enrichment. Localized accessibility surges within broader genomic compaction align with mammalian viral paradigms: lytic Epstein-Barr virus reactivation drives focal opening alongside global host closure (38), whereas early human cytomegalovirus infection suppresses de novo transcription without altering chromatin accessibility (39). Cross-system accessibility dynamics thus lack a unified directional trend. Diverging from the genomic baseline, outlier domains exhibited distinct physical behavior.

Nucleosome positioning across transcription start sites (TSSs) resolved regional chromatin responses (Fig. 6). In control populations, SOC and standard chromosomes exhibited maximal accessibility fluctuations between 0 and 10 h, tracking normal diel expression (Fig. 6, A to C). During infection, these sectors diverged sharply. Standard TSS accessibility increased marginally across infection (*P* < 0.05, Mann-Whitney *U* test; Fig. 6, D and G), whereas BOC1 and SOC1 TSSs experienced a rapid accessibility spike at 1 hpi before steadily waning (*P* < 0.0001, Mann-Whitney *U* test; Fig. 6, E, F, H, and I). This initial opening respected pre-existing domain boundaries: of 92 loci gaining accessibility at 1 hpi, 75 mapped directly within BOC1 or SOC (fig. S13E; data S16). Opening loci encompassed metabolic processes, nucleic acid machinery, structural components, transport systems, and signaling pathways, including RNA polymerase III subunits that displayed relative upregulation on BOC1 at 6 hpi. Early chromatin opening within outlier domains thus anticipated downstream transcriptional divergence by several hours, preceding both Hi-C sampling points.

**Fig. 6.**
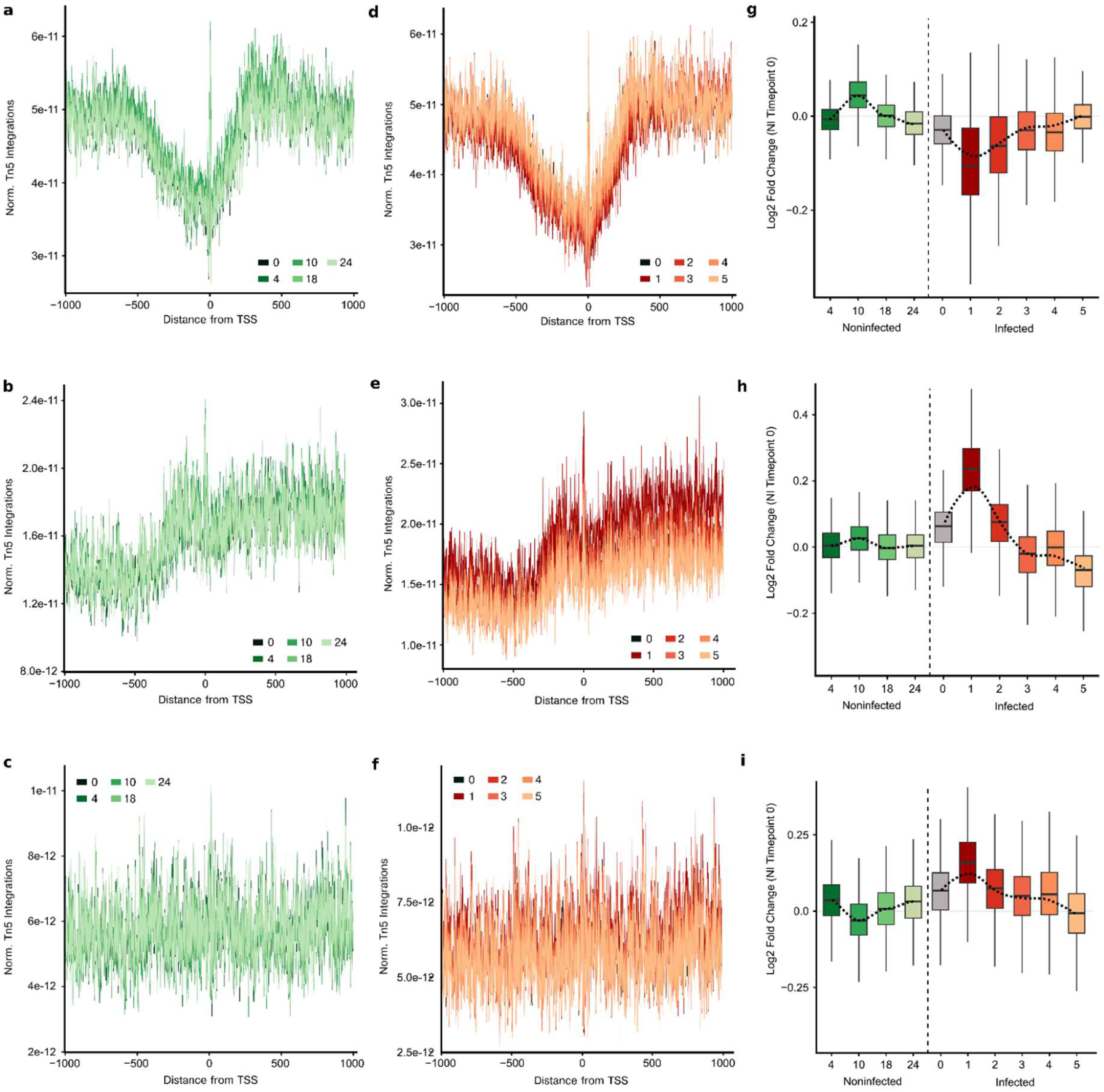
Region-specific chromatin accessibility dynamics during OtV5 infection. (**A** to **C**) Metaplots of sequencing-depth-normalised Tn5 integration (ATAC-seq) around transcription start sites (TSSs) in the standard nuclear region (A), BOC (chromosome 2; B), and SOC (chromosome 19; C) in noninfected cells sampled across the diel cycle (0, 4, 10, 18, 24 h). (**D** to **F**) As in (A) to (C) but for OtV5-infected cells sampled hourly through early infection (0 to 5 hpi): standard region (D), BOC (E), and SOC (F). (**G** to **I**) Distributions of log2 fold change in normalised Tn5 integration relative to the noninfected 0 h timepoint for the standard region (G), BOC (H), and SOC (I); the vertical dashed line separates noninfected (green) from infected (orange-red) samples, and the smoothed loess line shows the mean trend. The dip in Tn5 signal at the TSS corresponds to the nucleosome-depleted region.

Chromatin accessibility fails to account for viral depletion, requiring distinction between two key spatial scales. In absolute terms, outlier cores maintained open chromatin while recording the lowest viral contact density across the host genome; high viral contact intersected elevated accessibility exclusively at SOC telomeres at 2 hpi, mapping to repeat-dense, GC-standard regions outside the outlier boundary before subsiding by 6 hpi. Intra-regional rank correlations between accessibility and viral contact remained positive across standard chromosomes (mean ρ = 0.297 at 2 hpi; mean ρ = 0.165 at 6 hpi; *P* < 0.05) and within outlier domains, peaking on SOC (mean ρ = 0.948 at 2 hpi; mean ρ = 0.673 at 6 hpi; *P* < 0.05; nine-bin estimate provided as indicative; data S17). Local accessibility thus governs intra-domain viral positioning but fails to predict inter-domain contrasts, which strictly track architectural boundaries. Chromatin accessibility, the classical metric for viral targeting, is decoupled from viral DNA positioning within outlier compartments.

## Discussion

### Viral takeover follows a pre-existing genomic map

Infection by OtV5 achieves nearly total population clearance by 24 hpi alongside a global host transcriptional collapse that remains robust after standardising for cell density and viral transcript displacement. This per-cell normalization decouples true transcript loss from compositional artifacts driven by massive viral expression (28). However, host shutoff proceeds non-uniformly: host expression changes, spatial insulation erosion, early chromatin opening, and viral DNA positioning are all bounded by coordinates established prior to viral entry. Three independent assays, conducted across infection, converge precisely onto this pre-existing architectural baseline. Viral takeover is thus spatially partitioned along coordinates mapped before infection.

Concurrently, OtV5 executes a temporally orchestrated expression cascade. Viral loci segregated into early, mid, and late kinetic classes; early genes encode transcription initiation factors and DNA topology machinery, mid-phase loci govern nucleotide biosynthesis, and late loci express structural virion components, with early and late classes displaying distinct promoter motifs (33, 34, 40–42). Early expression of host-directed factors indicates active nuclear co-optation rather than immediate host genomic cleavage (13, 43), distinguishing OtV5 from chloroviruses that degrade host DNA within minutes via virion-packaged nucleases (44). Furthermore, early upregulation of host H2B and H3 histone variants aligns with chromatinisation of viral DNA within the host nucleus (13). OtV5 thus systematically reorganizes the host nuclear space rather than triggering immediate destruction, carrying out viral takeover through a spatially partitioned architecture.

### Compartment identity predicts viral DNA distribution

Within this structural framework, viral inter-genome contact profiles resolve two competing models that remain inextricably linked in animal models. Nuclear DNA viral episomes partition heterogeneously across host nuclear space along functional axes: active transcription units target accessible, expression-dense loci (15, 16), whereas latent genomes associate with AT-rich, silenced heterochromatic domains (2, 17–20). These target features are structurally confounded in animal hosts because repressive nuclear compartments are intrinsically AT-rich and transcriptionally silent. In *O. tauri*, however, sequence properties and expression dynamics dissociate. BOC1 and SOC1 mirror the primary sequence hallmarks of animal repressive compartments—exhibiting low GC content, transposon accumulation, and hypomethylation—yet represent the most transcriptionally resistant genomic sectors while undergoing rapid chromatin opening. Paradoxically, OtV5 DNA is systematically depleted from these loci, reaching minimum contact density across BOC1, while preferentially engaging accessible standard-GC telomeres at 2 hpi. Domain identity, rather than local accessibility or host transcriptional activity, dictates viral positioning. Crucially, whereas animal paradigms reflect persistent or latent infections (frequently involving integration-competent viruses), OtV5 executes rapid lytic takeover over hours, driving viral exclusion relative to pre-existing, spatially defined domain boundaries rather than broad chromosome-wide biases.

Nuclear DNA viruses often construct localized replication factories that marginalize host chromatin toward the nuclear periphery (45), parallel to phage-encoded nuclear shells that physically separate viral from host genomic material (46). Reduced viral-host contact could theoretically reflect physical occlusion by viral factories rather than domain-specific exclusion. Nevertheless, factory formation fails to explain internal regional contact contrasts within identical nuclei or the intense early targeting of SOC telomeres. Super-resolution spatial mapping of viral DNA relative to outlier domains will be needed to test this directly.

### Infection erodes pre-existing genome architecture

Multiple spatial observations confirm that pre-existing genome architecture, rather than infection-driven spatial reorganization, governs this response. First, the sharp expression breakpoint resolved on BOC at 6 hpi coincides precisely with the ∼535 kb insulation coordinate delineated in uninfected hosts (7). Second, the elevated trans-interaction frequency between BOC1 and SOC1 precedes viral entry and is maintained, albeit diminished, during takeover. Because physical proximity dictates contact probability (36), these outlier compartments reside in mutual nuclear proximity prior to viral arrival. Pre-existing spatial organization thus constitutes a baseline feature that dictates physical and functional outcomes during viral takeover.

These functional and structural datasets remain mechanistically intertwined. Active transcription elongation directly influences 3D chromatin conformation (47), whereas viral transcription read-through during herpesvirus infection evicts structural proteins and opens host loci (24). Consequently, retained expression within BOC1 and SOC1 may actively drive local insulation decay rather than simply coinciding with structural changes. This multi-omic alignment maps the physical boundaries of host takeover, though distinct molecular pathways may operate within these shared coordinates.

Rather than remaining static, this architectural framework undergoes progressive breakdown. Outlier regions exhibit selective erosion of spatial insulation, registering structural disruption early despite maintaining relative transcriptional output. Interchromosomal trans contacts between BOC1 and SOC1 fall to 0.65 of baseline by 6 hpi, while regional chromatin accessibility surges at 1 hpi before steadily waning. Concurrently, total cellular transcript pools drop 9-fold to 18-fold across all sectors. These coupled dynamics reveal an ordered cascade, beginning with early accessibility spikes, progressing through domain-restricted transcriptional retention, and culminating in the movement of viral DNA from SOC telomeres toward chromosomes 17 and 20.

### BOC1 and SOC1 respond along different axes

The genomic sectors separated by this structural boundary exhibit functional asymmetry. BOC1 represents a primary structural anomaly, recording minimal viral DNA interaction, exclusive coupling between expression and local physical contacts, and strict confinement of its transcriptional response within a sharp coordinate. However, total transcript yield from BOC1 decayed rapidly alongside standard chromosomes (log2 -4.1 versus -4.2 at 10 hpi), accompanied by a declining regional output share. Conversely, SOC1 forms a distinct expression sanctuary, progressively expanding its overall transcript contribution throughout infection while preserving double the per-cell transcript density of standard chromatin and demonstrating strong inverse coupling between viral engagement and expression. Outlier chromosome 19 was previously termed a viral immunity chromosome due to concentrated structural variations and expression alterations during host resistance (48–50), with resistant *O. mediterraneus* lineages accumulating translocations and massive deletions within SOC (51). While our experiments examine lytic takeover rather than immunity, these findings demonstrate that chromosome 19 mounts a specialized physical and transcriptional program during active viral attack. Spatial confinement also differs between domains: structural impacts remain strictly internal to BOC1, whereas SOC1 transcriptional dynamics spill over into adjacent telomeric flanks. Paradoxically, these same telomeres recorded peak viral contact density at 2 hpi. Consequently, two domains sharing identical nuclear positioning, sequence profiles, and spatial insulation respond to infection along divergent functional axes, requiring future models to account for this inherent asymmetry.

These patterns could alternatively reflect pre-existing chromatin composition. Both BOC1 and SOC1 feature low-GC composition, transposon enrichment, and hypomethylation (4, 5), features known to drive nuclear compartmentalization in eukaryotic genomes. These characteristics can decouple, as seen in *Bathycoccus* where outlier chromosomes persist without transposable element accumulation (9). Spatial co-localization, domain insulation, and expression retention might therefore stem entirely from constitutive compartment identity established prior to viral entry (7). Under this model, sequence composition establishes the structural architecture governing host takeover. Crucially, pre-existing compartment identity fails to explain. the local depletion of viral DNA, a physical exclusion phenomenon without parallel in viral systems Resolving these models will require direct epigenetic manipulation of host outlier chromatin prior to viral infection.

### Genome organisation structures both sides of the interaction

Host and viral genomes thus function as two interlinked, parallel maps. Along the host genome, pre-existing spatial boundaries predict where lytic takeover is most strongly resisted. Reciprocally, the OtV5 locus exhibiting maximal host contact at 2 hpi overlaps the hypervariable genomic window known to diversify rapidly across isolates and govern host range (37), aligning host specificity directly with host physical engagement. Although this contact enrichment is subtle relative to a permutation baseline, positional organization structures both genomes in advance. Mechanistically, this parallelism holds: host engagement tracks compartment identity rather than active transcription, while the host-targeted viral locus encodes late structural factors that remain transcriptionally silent during early infection. Host-viral spatial interactions are thus decoupled from active transcription across both genomes.

Distinctive genomic compartments occur widely across eukaryotic lineages. Fungal accessory chromosomes form repeat-dense, structurally dynamic domains maintained under unique chromatin states (53–55), whereas brown algal sex chromosomes concentrate transposable elements and novel loci within non-recombining regions (56, 57). What sets *O. tauri* apart is the dynamic behavior of its outlier domains during lytic attack. While host chromatin boundaries constrain viral interactions in select mammalian systems (27), we demonstrate here that pre-existing spatial architecture partitions the host transcriptional shutdown, insulation breakdown, and chromatin opening during productive viral takeover in a microbial eukaryote. Furthermore, this work provides the first evidence of viral DNA depletion from defined host compartments, as well as the initial high-resolution profiling of chromatin accessibility dynamics across infection in a marine picoeukaryote.

### Conclusions

Disentangling infection-driven transcriptional programs from constitutive chromatin architecture requires targeted functional perturbations of host outlier domains prior to viral challenge to test if viral exclusion persists. Because interchromosomal trans interactions between BOC1 and SOC1 are inherently symmetric, physical proximity alone cannot establish functional coordination; targeted deletion or epigenetic manipulation of a single outlier domain will determine whether its partner responds interdependently or merely shares adjacent nuclear space. Higher-resolution sampling time courses combined with reduced multiplicity of infection will further establish the kinetic bounds of this structural partitioning. Finally, extending these parallel multi-omic profiles to related Mamiellales genera and diverse marine eukaryotic lineages will reveal whether pre-existing nuclear architecture broadly structures lytic viral takeover across ocean ecosystems or represents an evolutionary feature unique to *O. tauri*.

In *O. tauri*, lytic viral takeover progresses heterogeneously across host nuclear space, organized by pre-established architectural coordinates rather than emergent spatial features. Pre-existing domain boundaries delineate the regional host response across transcriptional retention, insulation decay, chromatin opening, and inter-genomic contact density. Nuclear genome architecture does not merely serve as a passive background during infection, but actively partitions the physical and functional takeover of the host cell.

## Materials and methods

### Algal culture

*O. tauri* strain RCC4221 was maintained in L1 medium (58) in non-treated polycarbonate flasks in a 14 h:10 h light:dark cycle at 35 μmol m-2 s-1 irradiance and 20 °C. To achieve a near-axenic culture, we isolated cells by flow cytometric cell sorting (BD Influx, Becton Dickinson) and maintained algal cells in exponential growth by weekly subculturing. A monthly antibiotic treatment prevented bacterial overgrowth (5 μg/mL ampicillin, 1 μg/mL gentamicin, 2 μg/mL kanamycin, 10 μg/mL neomycin, final concentrations). Algal and bacterial growth was regularly monitored by flow cytometry (Cytoflex, Beckman Coulter) based on side-angle light scatter and chlorophyll red fluorescence; bacterial cells were quantified after SYBR Green I staining.

### Virus production

The virus used was OtV5 (NCBI accession no. NC_010191.2). To infect large volumes at the high multiplicity of infection (MOI) needed to ensure that a large majority of cells were infected, we successively upscaled virus stocks. We began by infecting small volumes (20 mL) of *O. tauri* cultures in exponential growth, allowing 3 days for complete lysis. Lysates were harvested and centrifuged at 4000*g* for 30 min to clear cell debris, and cleared lysates were used to initiate larger volume (> 200 mL) infections. Stocks were upscaled from 15 mL to 40 L over several months. Immediately before the final experiment, stocks were concentrated to 500 mL using a VivaFlow concentration system (Sartorius, Germany) with a 30 kDa PES membrane.

### Viral enumeration

Plaque-forming units for concentrated OtV5 lysate were calculated by plaque assay (11). For virus-like particles (VLPs) per mL in concentrated lysate and in *O. tauri* samples, 1 mL was fixed with glutaraldehyde (0.5% final concentration), incubated in the dark for 15 min at 4 °C, flash-frozen in liquid nitrogen, and stored at -80 °C. Defrosted samples were diluted in TE buffer (10 mM Tris-HCl, 1 mM EDTA, pH 8) and stained with SYBR Green at a final concentration of 5 × 10-5 commercial stock (Invitrogen S7585), incubated for 10 min at 80 °C then 5 min at room temperature. Samples were run on a FACScan flow cytometer (Becton Dickinson) triggered on green fluorescence at low flow (approximately 20 μL min-1) for 1 min, with event rates between 100 and 500 cells s-1 (59).

### Viral infection

We grew 18 L of *O. tauri* in a large glass Duran bottle, with no antibiotics in the 3 weeks preceding the experiment. The stock culture was split into eight flasks (2.2 L each), four controls (A to D) and four for infection (E to H). Infection flasks were inoculated with 60 mL of concentrated OtV5 lysate; controls received 60 mL of L1 medium. Based on quantification of the virus stock, the MOI was 9 by flow cytometry and 7 by plaque assay. The experiment was initiated 1 h into the light cycle and sampled at 2 h intervals for the first 14 h, followed by a 4 h interval (18 h) and then a 6 h interval (24 h). Sampling times were adjusted as the experiment progressed because cell lysis began earlier than expected from a previous publication (13). At each sampling point, flasks were gently mixed and the total volume required for all measurements was decanted into a glass vessel for subsampling.

### Additional infection for ATAC-seq

For the small-scale infection experiment, we grew 0.5 L of *O. tauri* in a glass conical flask, with all work carried out in a microbiological cabinet. The stock culture was split into six flasks (approximately 60 mL each), three controls and three for infection. Each infection flask was spiked with 3.75 mL of concentrated OtV5; an equivalent volume of 0.1 μm filtered L1 medium was added to the controls. Flasks were shaken throughout using an electronic shaker. Quantification of the virus stock by flow cytometry gave an MOI of approximately 7. The experiment was initiated 0.5 h into the light cycle and sampled at 1 h intervals for the first 6 h, followed by timepoints representing cell-cycle phases.

### Scanning electron microscopy

Cells in culture were fixed in 2% glutaraldehyde and 2% paraformaldehyde in 0.1 M sodium cacodylate buffer, pH 7.2, for 1 h at room temperature, washed 3 × 5 min in buffer, and post-fixed in 1% aqueous osmium tetroxide for 1 h. After 3 × 5 min washes in deionised water, cells were dehydrated in a graded ethanol series from 30% through 95% (5 min per step), then 2 × 100% for 10 min each. Cells were passed through a polycarbonate filter with a 200 nm pore size under mild vacuum. Filters with adhered cells were incubated for 3 min in hexamethyldisilazane (Merck, Gillingham, UK) and air-dried before mounting on aluminium stubs using double-sided carbon tabs. Samples were coated with 10 nm gold/palladium (80:20; Q150T sputter coater, Quorum, Laughton, UK) before imaging with a Zeiss Gemini500 SEM operated at 5 kV with a secondary electron detector (SE2).

### Transmission electron microscopy

Cells were processed as described for SEM, except that post-fixation used 1% osmium reduced in 1.5% potassium ferrocyanide in cacodylate buffer. Following gradual dehydration as described above, cells were infiltrated with Durcupan resin (Sigma-Aldrich, Gillingham, UK) by incubating overnight in a 1:1 mix of 100% ethanol and resin and the following day twice in fresh resin for a total of 8 h. Samples were placed in BEEM capsules and polymerised at 62 °C for 24 h. Ultrathin (70 nm) sections were collected on pioloform-coated copper EM grids and contrasted using Reynolds lead citrate. Images were taken using a JEOL JEM 1400 transmission electron microscope operated at 120 kV with a digital camera (ES1000W, Gatan, Ametek, Leicester, UK).

### RNA extraction and sequencing

*O. tauri* RCC4221 cells were collected by centrifuging a 20 mL sample (10 min at 4000*g* at 4 °C), snap-frozen in liquid nitrogen, and stored at -80 °C. RNA was extracted using an RNeasy Plant kit (QIAGEN) with RLT buffer incubated at 56 °C for 3 min. Residual DNA was digested using the Turbo DNA-free kit (Thermo Fisher Scientific). RNA was quantified using a Qubit fluorometer (Thermo Fisher Scientific), and quality was assessed using an Agilent 2100 Bioanalyzer system. Extracts with an RNA integrity number greater than eight were subjected to poly(A) selection prior to library preparation with the Illumina TruSeq Directional Library Prep Kit and sequenced on an Illumina NovaSeq 6000 (150 bp paired-end reads).

### RNA-seq data processing

Adapters and low-quality reads were trimmed using fastp v0.23.2 (60) (minimum length 50). Quality-control reports were compiled using FastQC v0.11.9 (61), MultiQC v1.13 (62), and fastp. Ribosomal RNA reads were removed using SortMeRNA v4.3.6 (63) against rRNA sequences from the SILVA database (64). Remaining reads were aligned and quantified to the *O. tauri* reference genome (GCF_000214015.3) using STAR v2.7.10 (65) with default parameters. Outlier-region boundary coordinates were verified by re-mapping against the telomere-to-telomere assembly of *O. tauri* RCC4221 (8), which resolves previously collapsed duplicated sequence within both outlier chromosomes; coordinates are reported on GCF_000214015.3 for comparability with earlier work. Counts were divided by gene length to standardise transcript levels for descriptive analyses; raw counts were used for differential expression. Differential expression between noninfected and infected samples and between timepoints was analysed with DESeq2 (35) (|log2FC| > 1, adjusted *P* < 0.1, design = ∼condition). Dimension reduction used vegan v2.6-4 (66) and FactoMineR v2.6 (67) (metaMDS and PCA). Non-negative matrix factorisation was used to cluster sample transcription and identify genes unique to each cluster, using the R package NMF v0.30.4.900 (68). NMF decomposes the expression count matrix into a product of two matrices.

### Per-cell normalisation of transcript abundance

Because differential expression and relative abundance measures are compositional, and because both cell number and viral transcript load change during infection, we derived an approximate per-cell measure of host transcript abundance. For each timepoint and condition, mapped host reads were divided by the number of cells contributing to the extraction, calculated as the flow-cytometric cell density multiplied by the 20 mL sample volume, and expressed as mapped host reads per 10^6^ cells (fig. S1). Conditions were compared by Welch’s *t* test on the derived per-cell means with *n* = 4 biological replicates, with Benjamini-Hochberg correction across timepoints.

To assess whether the reduction in host reads was explained by viral transcripts occupying a pool of fixed size, expected host reads in the infected library were calculated as the noninfected host read count multiplied by the host fraction of the infected library, and compared with the observed count. This follows established practice for transcriptomes undergoing global change, where compositional normalisation misreports per-cell output (28–31). It removes the cell-number confound but not the effect of library-size normalisation during library preparation and sequencing, for which exogenous spike-in controls would be required. Values are therefore interpreted as bounds on the magnitude of the transcriptional collapse rather than as absolute transcript counts.

### Hi-C sample collection and sequencing

Crosslinking was performed by adding formaldehyde (2% v/v final concentration; 16% TEM grade, Sigma) directly to the culture medium containing cells at targeted timepoints (60 mL) and incubating at room temperature for 20 min. Crosslinking was quenched with glycine (0.2 M final concentration) for 5 min at room temperature on a roller. Cells were washed with ice-cold PBS and stored at -80 °C. The final 12 Hi-C libraries, covering triplicate 2 hpi and 6 hpi control and infected samples, were generated using the Proximo Microbe Hi-C kit v2 (Phase Genomics, USA) following the manufacturer’s instructions. Libraries were quality-assessed for size by electrophoresis and for amplification efficiency by qPCR, and sequenced on the Illumina NovaSeq 6000 platform.

### Hi-C data processing

Adapters and low-quality reads were trimmed using fastp v0.23.2, and quality-control reports were compiled with FastQC v0.11.9, MultiQC v1.13, and fastp. Samples were pooled before running the Hi-C processing pipeline (restriction sites “GATC,AATT”; ligation sites “GATCGATC,GATCAATT,AATTGATC,AATTAATT”), based on Nextflow v22.04.5 (nf-core/hic v1.3.0, available as a Docker profile) (69–71). Reads were mapped to the *O. tauri* reference genome (GCF_000214015.3) using Bowtie 2 v2.3.5.1 (72). Duplicate removal and contact-pair validation used HiC-Pro v2.11.1 (71). Raw and normalised contact maps were generated using cooltools v0.3.0 (73) at bin resolutions of 5, 10, and 20 kb. Additional normalisation used “iced” with HiC-Pro, a Python implementation of the iterative correction algorithm (74). Hi-C quality reports were generated using HiCUP (75). Samples were pooled by replicate and Spearman correlations calculated on cool files (fig. S8).

Insulation scores were computed with cooltools (52) on balanced 10 kb matrices using a 50 kb sliding diagonal window and reported as log2 values. Under this definition a low score indicates few contacts crossing the window and therefore strong insulation, and domain boundaries correspond to local minima. Change in insulation (Δ) is reported as the infected value minus the noninfected value for each 10 kb bin. Genomic bins were assigned to Standard, BOC1, nonBOC1, or SOC1 by chromosome and coordinate, and each annotated region was compared with the Standard background within each sample by two-sided Mann-Whitney *U* test with FDR correction. Raw insulation scores by region and timepoint are given in fig. S11. Compartment eigenvector decomposition and contact-frequency derivatives were computed with cooltools; the eigenvector result is reported in fig. S16 and data S9.

Interchromosomal contact frequencies were extracted from balanced matrices and grouped into four classes: BOC-SOC (1 pair), BOC-standard (18 pairs), SOC-standard (18 pairs), and standard-standard (153 pairs). Because the BOC-SOC class contains a single region pair, this comparison is reported descriptively.

Host-virus contacts were extracted from balanced matrices at 10 kb resolution at 2 and 6 hpi and compared between annotated regions and the Standard background by two-sided Mann-Whitney *U* test with FDR correction. Spearman correlations between contact frequency and gene expression were computed across gene loci within each region.

Mann-Whitney tests applied to genomic bins treat spatially autocorrelated observations as independent and therefore return anticonservative *P* values; reported significance should be read as an effect-size indicator supported by the direction and consistency of the regional differences. Replicate-aware methods for differential Hi-C analysis (76) were not applicable here because libraries were pooled before processing.

Host-virus contact along the OtV5 genome was summarised as a moving average in 20 kb windows (fig. S13; data S14) and local maxima identified from the smoothed profile. Enrichment over the FhaB gene block (OtV5_070/OtV5_071; 54,743-62,876 bp) and over a 20 kb sub-window (45,000-65,000 bp) of the hypervariable region (25,000-80,000 bp) defined across O. tauri-infecting viruses (37) was expressed as the ratio of mean contact inside the window to that outside it, computed on balanced 5 kb matrices with host contacts summed across chromosomes 1 to 20 and organelle contigs excluded. Significance was assessed by permuting a width-matched window to 10,000 random positions along the viral genome (empirical P = [number of null ratios greater than or equal to the observed + 1] / 10,001).

### ATAC sampling, extraction, and sequencing

*O. tauri* RCC4221 cells were collected by taking 0.5 mL of culture, adding DMSO (10% final concentration), and placing samples in a Mr. Frosty freezing container (Nalgene) at -80 °C. ATAC experiments were performed on 100,000 cells from thawed samples according to the manufacturer’s protocol (Diagenode cat. no. C01080001). PBS washes in this protocol were replaced with L1 medium washes to prevent early osmotic shock. DNA was quantified using a Qubit fluorometer (Thermo Fisher Scientific), and quality was assessed on an Agilent 2100 Bioanalyzer before sequencing on an Illumina NovaSeq X (150 bp paired-end reads).

### ATAC-seq data processing and analysis

Raw paired-end reads from 10 biological conditions, each with three replicates, were processed using a standardised pipeline. Adapter trimming and quality filtering used fastp v0.23.2 (minimum Phred score 20, minimum read length 30 bp). Trimmed reads were aligned to the reference genome using Bowtie 2 v2.4.5 (72) with the --very-sensitive preset and a maximum fragment size of 2000 bp. Aligned reads were converted to BAM format, sorted, and PCR duplicates removed using samtools (77). To account for Tn5 transposase insertion offset, deduplicated BAM files were corrected using the --ATACshift function of deepTools alignmentSieve (78). Peak calling was performed per replicate using MACS2 v2.2.7.1 (79) with paired-end BAM input, genome size adjusted for the species, and --call-summits enabled. Cross-sample peak comparison used DiffBind v3.20.0 (80, 81) with edgeR (82). Footprinting and nucleosome-positioning analyses were performed in R using an existing script (83) with VplotR (84) and rtracklayer (85). Merged paired-end BAM files were analysed against genomic regions provided as BED files. Tn5 insertion footprints were computed with VplotR using strand-agnostic, centrally aligned aggregation across multiple window sizes (±50 bp, ±100 bp, ±1000 bp), normalised by library depth. Statistical differences in accessibility within ±25 bp of TSS centres were assessed using paired and unpaired *t* tests across conditions. V-plots were generated to visualise fragment-length distributions relative to TSS centres.

### Motif detection

MEME was used in Differential Enrichment mode, with target and control sequences representing the promoters of the genes in the regions of interest (86). Promoters were extracted using Python, taking 500 bp upstream and 100 bp downstream of the putative TSS, interchanging these values for genes on the negative strand. A second sequence set represented the intergenic regions of interest.

### Homology, GO, and other analyses

Fisher’s exact test (one-tailed) was performed using a custom Python script on gene subsets to compare enriched terms over target regions versus the rest of the genome. Signal peptides were identified within upregulated proteins using SignalP v6.0 (87). AlphaFold structure prediction was used to annotate viral proteins of unknown function, for example the UBOX-domain identification for OtV5_154c; updated annotations were determined using AlphaFold3 (88) and Foldseek (89).

### Statistical analysis

Unless stated otherwise, comparisons between annotated genomic regions and the Standard background used two-sided Mann-Whitney *U* tests with Benjamini-Hochberg FDR correction, and correlations were Spearman rank coefficients tested against zero. Per-cell transcript comparisons used Welch’s *t* test with Benjamini-Hochberg correction. Regional expression shares were compared by Wilcoxon signed-rank test across timepoints. Viral-host contact enrichment over the FhaB block and the hypervariable window was assessed by width-matched permutation along the viral genome (10,000 permutations). Significance thresholds are given in each figure legend. RNA-seq and flow cytometry used *n* = 4 biological replicates; ATAC-seq used *n* = 3; Hi-C libraries were prepared in triplicate and pooled before processing, so Hi-C comparisons are made across genomic bins rather than across biological replicates.

## Supplementary Materials

Supporting Information Text (Extended Methods) Figs. S1 to S13 Tables S1 to S3 Data S1 to S17 SI References

## Acknowledgments

We are grateful to Karen Moore and Bryony Williams (Exeter Sequencing Facility, University of Exeter) for their technical advice on Hi-C and ATAC sequencing. We thank Christian Hacker (Bioimaging Centre, University of Exeter) for providing the TEM and SEM data, and Aurélie Chambouvet and Evelyne Derelle (CNRS, France) for providing algal and virus strains. We thank Ashley Bell, Victoria L. N. Jackson, and Sara Castillo Vila for helping to sample timepoints, Margot Tragin for preliminary RNA-seq analyses, and Bryony Williams and Mike Allen for feedback on the manuscript.

## Author contributions

Conceptualization: A.M.; Methodology: K.H., M.V., S.K., A.M.; Investigation: K.H., M.V., E.S.K., S.K.; Formal analysis: K.H., Y.L.-M., R.R.; Software: K.H., Y.L.-M., R.R.; Visualization: K.H.; Supervision: A.M.; Funding acquisition: A.M.; Writing - original draft: K.H., A.M.; Writing - review and editing: all authors.

## Code availability

Custom scripts for promoter extraction, Fisher enrichment testing, per-cell normalisation, and host-virus contact profiling are deposited in the Zenodo repositories listed in the Data and materials availability statement.

## Data and materials availability

ATAC-seq, Hi-C, and RNA-seq reads are available in the ENA Sequence Read Archive under BioProjects PRJEB107987 and PRJEB107794. Additional data are available in Zenodo repositories (10.5281/zenodo.18938192). All data needed to evaluate the conclusions in the paper are present in the paper or the Supplementary Materials.

