## Supplementary Information for "Pre-existing genome architecture partitions viral takeover in the smallest free-living eukaryote"

**Short title:** Genome architecture partitions viral takeover

Keith Harrison[1,2], Martha Valiadi[1,2,3], Estelle S. Kilias[1,2,4], Yann Loe-Mie[5], Ramy Ragheb[1,2], Susan Kimmance[1,2] & Adam Monier[1,2*]

[1]Living Systems Institute, University of Exeter, Exeter, UK. [2]Biosciences, Faculty of Health and Life Sciences, University of Exeter, Exeter, UK. [3]Foundation for Research and Technology Hellas, Institute of Molecular Biology and Biotechnology, Heraklion, Crete, Greece. [4]Department of Biology, University of Oxford, Oxford, UK. [5]Institut Pasteur, Université Paris Cité, Bioinformatics and Biostatistics Hub, Paris, France.

**Competing Interest Statement:** No competing interests

**Keywords:** Chromosome conformation capture, picoeukaryotes, virus infection, transcriptomics.

**This PDF file includes:**

Figures S1 to S13

Tables S1 to S3

Legends for Datasets S1 to S17

SI References

**Other supporting materials for this manuscript include the following:**

Datasets S1 to S17

**Figures**

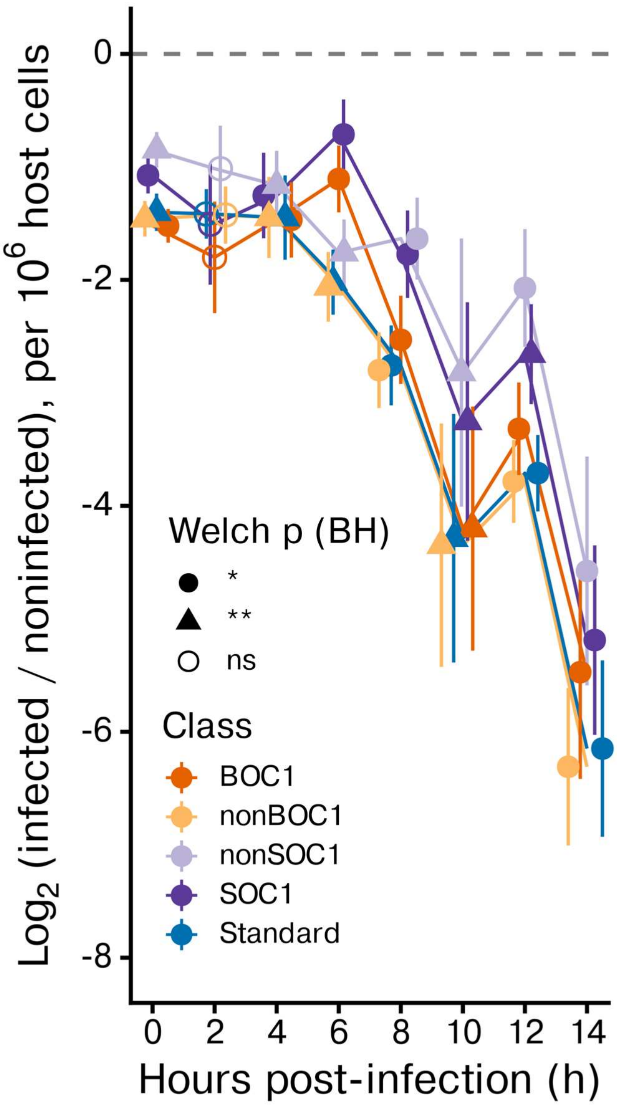


**Figure S1. Flow cytometry-normalized RNA-seq comparison between infected and noninfected samples.** $Log_2$ fold change of flow-cytometry host count-normalized mean transcript abundance between infected and noninfected samples across different transcript classes. Shape and fill represent Welch t-test FDR (BH) between mean transcript abundance normalized by flow cytometry host counts of infected and noninfected mean transcript abundances.

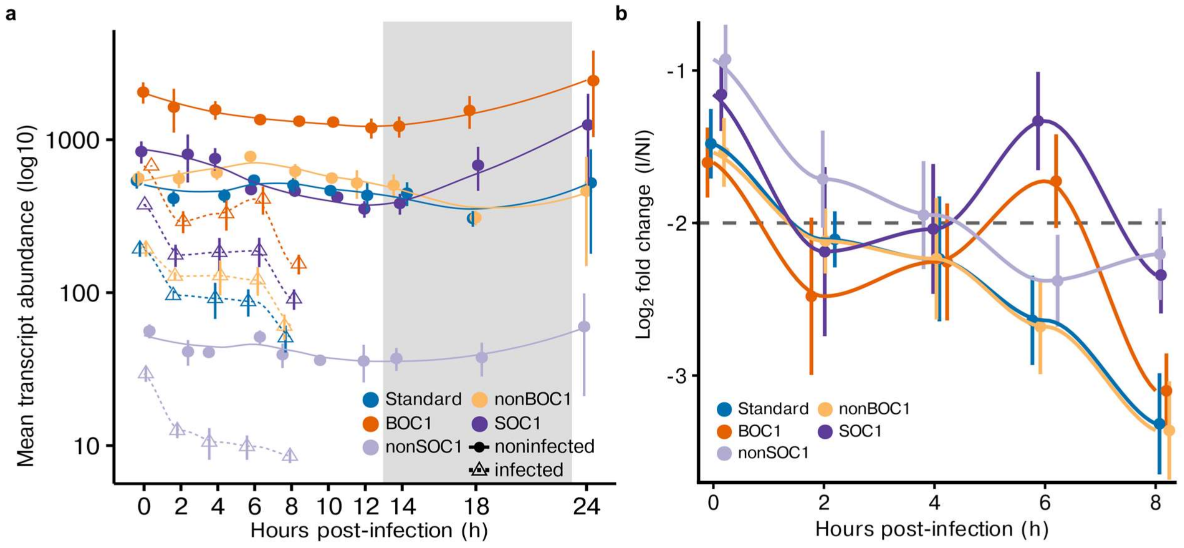


**Figure S2. Mean transcript abundance across each genomic region during OtV5 infection and diel cycle.** (**a**), Mean transcript abundance $log_{10}$ for noninfected and infected transcriptomes of *O. tauri*, genes are grouped based on encoding region: Standard, BOC1 (chr2 outlier region), nonBOC1 (chr2 non-outlier region), SOC1 (chr19 outlier region), nonSOC1 (chr19 non-outlier region). (**b**), Fold-change ($log_2$ ratio of infected to noninfected) in *O. tauri* transcript abundance over time. Genes are grouped based on encoding region: Standard, BOC1 (chr2 outlier region), SOC1 (chr19 outlier region), alongside non-outlier regions of BOC and SOC.

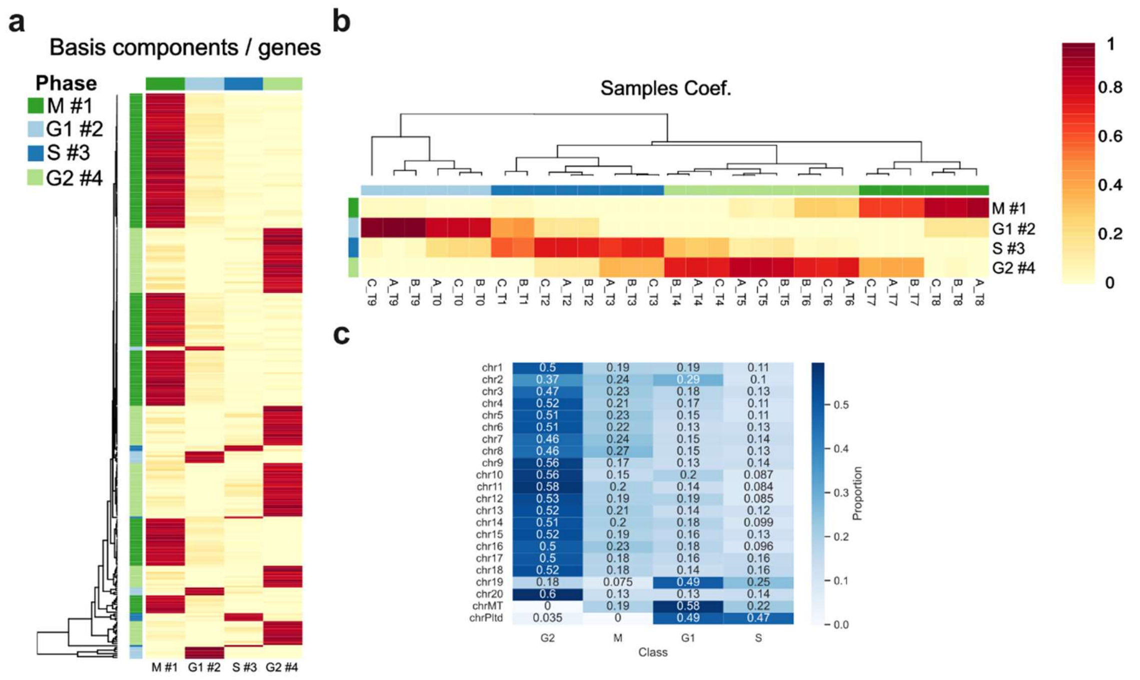


**Figure S3. Nonnegative matrix factorization (NMF) of noninfected *O.* tauri.** (**a**), Heatmap of basis components, genes and their correlation with each cluster. (**b**), Heatmap of the NMF coefficients matrix with cells coloured according to the correlation coefficients of each replicate's timepoint to each cluster. (**c**), Proportion of genes found in each cluster and the chromosome there are encoded on.

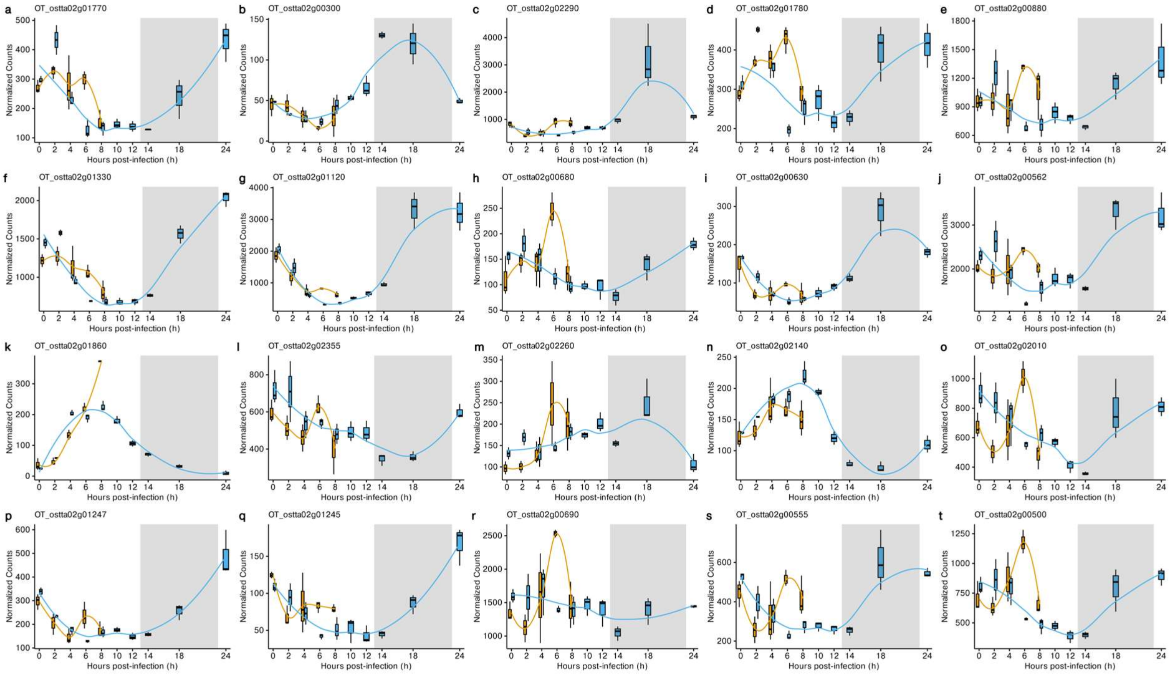


**Figure S4. Transcription of BOC1 DNA/RNA functionality-related genes.** (**a–l**), Normalised counts of transcript levels of genes conserved on BOC1 in Mamiellales with DNA/RNA functionality across infection in Noninfected (blue) and OtV5-infected (orange) *O. tauri* cells. (**a**), OT_ostta02g01770 (DNA-directed RNA polymerase subunit). (**b**), OT_ostta02g00300 (DNA-directed RNA polymerases I, II, and III subunit RPABC2). (**c**), OT_ostta02g02290 (DNA-directed RNA polymerase subunit beta). (**d**), OT_ostta02g01780 (RNA polymerase archaeal subunit P/eukaryotic subunit RPC10). (**e**), OT_ostta02g00880 (Plastid-encoded RNA polymerase subunit alpha). (**f**), OT_ostta02g01330 (H/ACA ribonucleoprotein complex subunit). (**g**), OT_ostta02g01120 (MAK16 homolog). (**h**), OT_ostta02g00680 (rRNA biogenesis protein RRP36). (**i**), OT_ostta02g00630 (Nucleotide-binding, alpha-beta plait). (**j**), OT_ostta02g00562 (Eukaryotic translation initiation factor 6). (**k**), OT_ostta02g01860 (Sister chromatid cohesion protein). (**l**), OT_ostta02g02355 (Transposase, Tc1-like). (**m**), OT_ostta02g02260 (Homeobox domain). (**n**), OT_ostta02g02140 (Zinc finger, PHD-finger). (**o**), OT_ostta02g02010. (**p**), OT_ostta02g01247 (Transposase, Tc1-like). (**q**), OT_ostta02g01245 (Transposase, Tc1-like). (**r**), OT_ostta02g00690 (DNA replication licensing factor MCM3). (**s**), OT_ostta02g00555. (**t**), OT_ostta02g00500 (P-loop containing nucleoside triphosphate hydrolase).

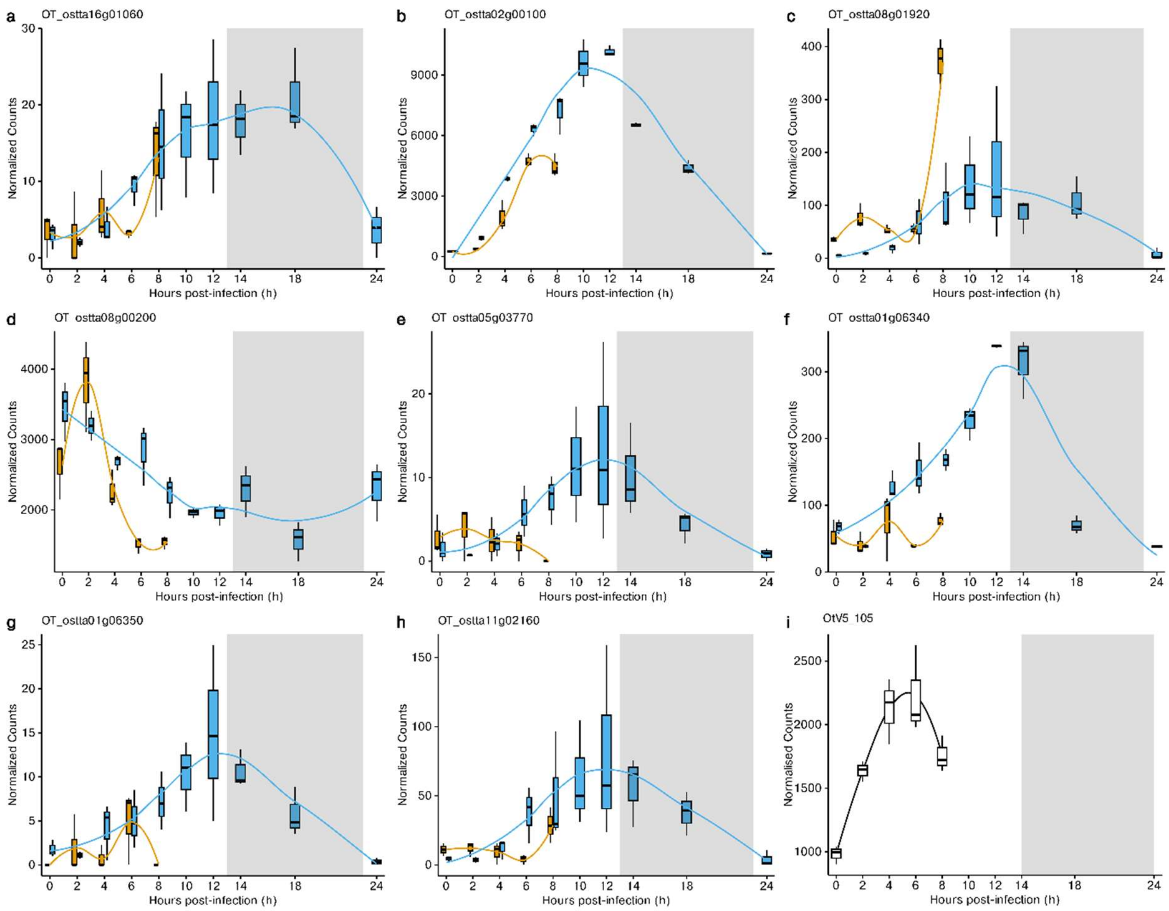


**Figure S5. Transcriptional profile of histone proteins within noninfected and OtV5-infected *O. tauri*.** (**a**), Histone H2A. (**b**), Histone H2A. (**c**), Histone H2B. (**d**), Histone H3. (**e**), Histone H3. (**f**), Histone H4. (**g**), Histone H4. (**h**), Histone H4. (**i**), Putative PCNA1_PBCV1 proliferating cell nuclear antigen-like 1.

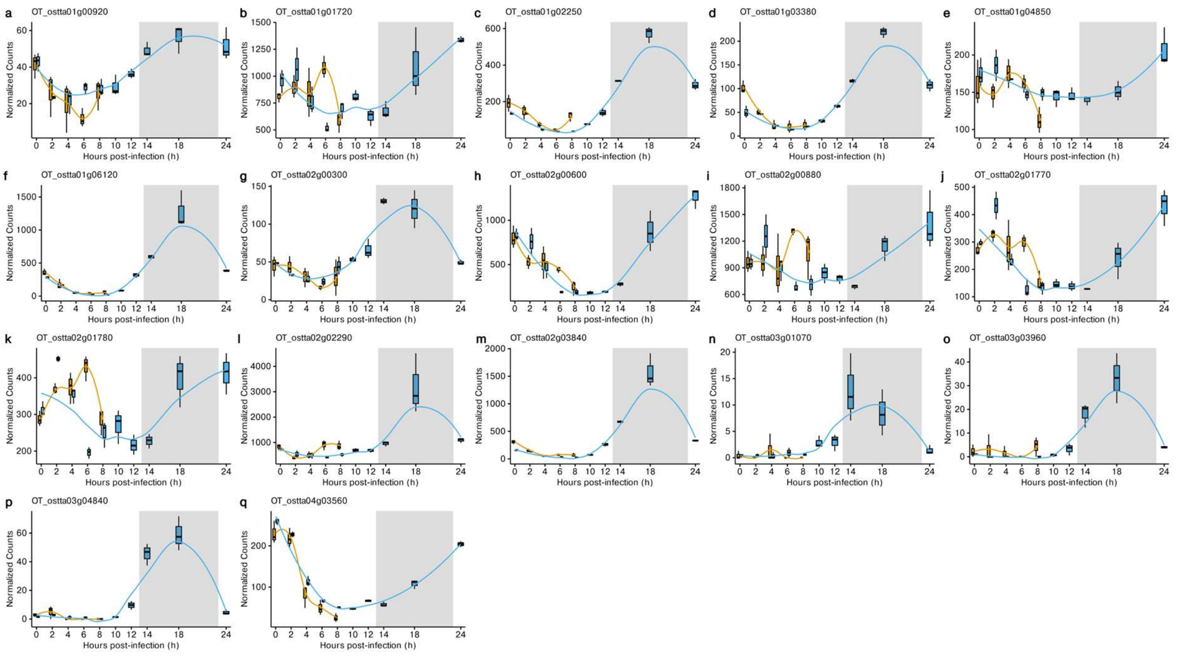


**Figure S6. Transcript dynamics of RNA polymerase genes across *O.* tauri during OtV5 infection.** (**a–l**), Normalised counts of transcript levels of RNA polymerase genes across *O. tauri* genome across infection in Noninfected (blue) and OtV5-infected (orange) *O. tauri* cells. (**a**), OT_ostta01g00920 (DNA-directed RNA polymerase, RBP11-like dimerisation domain). (**b**), OT_ostta01g01720 (DNA-directed RNA polymerase subunit beta). (**c**), OT_ostta01g02250 (DNA-directed RNA polymerase III subunit Rpc5). (**d**), OT_ostta01g03380 (RNA polymerase III Rpc4). (**e**), OT_ostta01g04850 (RNA polymerase II-associated, Paf1). (**f**), OT_ostta01g06120 (DNA-directed RNA polymerase subunit beta). (**g**), OT_ostta02g00300 (DNA-directed RNA polymerases I, II, and III subunit RPABC2). (**h**), OT_ostta02g00600 (DNA-directed RNA polymerase III subunit RPC3). (**i**), OT_ostta02g00880 (Plastid-encoded RNA polymerase subunit alpha). (j), OT_ostta02g01770 (DNA-directed RNA polymerase subunit). (**k**), OT_ostta02g01780 (RNA polymerase archaeal subunit P/eukaryotic subunit RPC10). (**l**), OT_ostta02g02290 (DNA-directed RNA polymerase subunit beta). (**m**), OT_ostta02g03840 (DNA-directed RNA polymerase subunit). (**n**), OT_ostta03g01070 (DNA-directed RNA polymerase Rpb11). (**o**), OT_ostta03g03960 (DNA-directed RNA polymerase subunit). (**p**), OT_ostta03g04840 (DNA-directed RNA polymerase subunit). (**q**), OT_ostta04g03560 (RNA polymerase sigma-70).

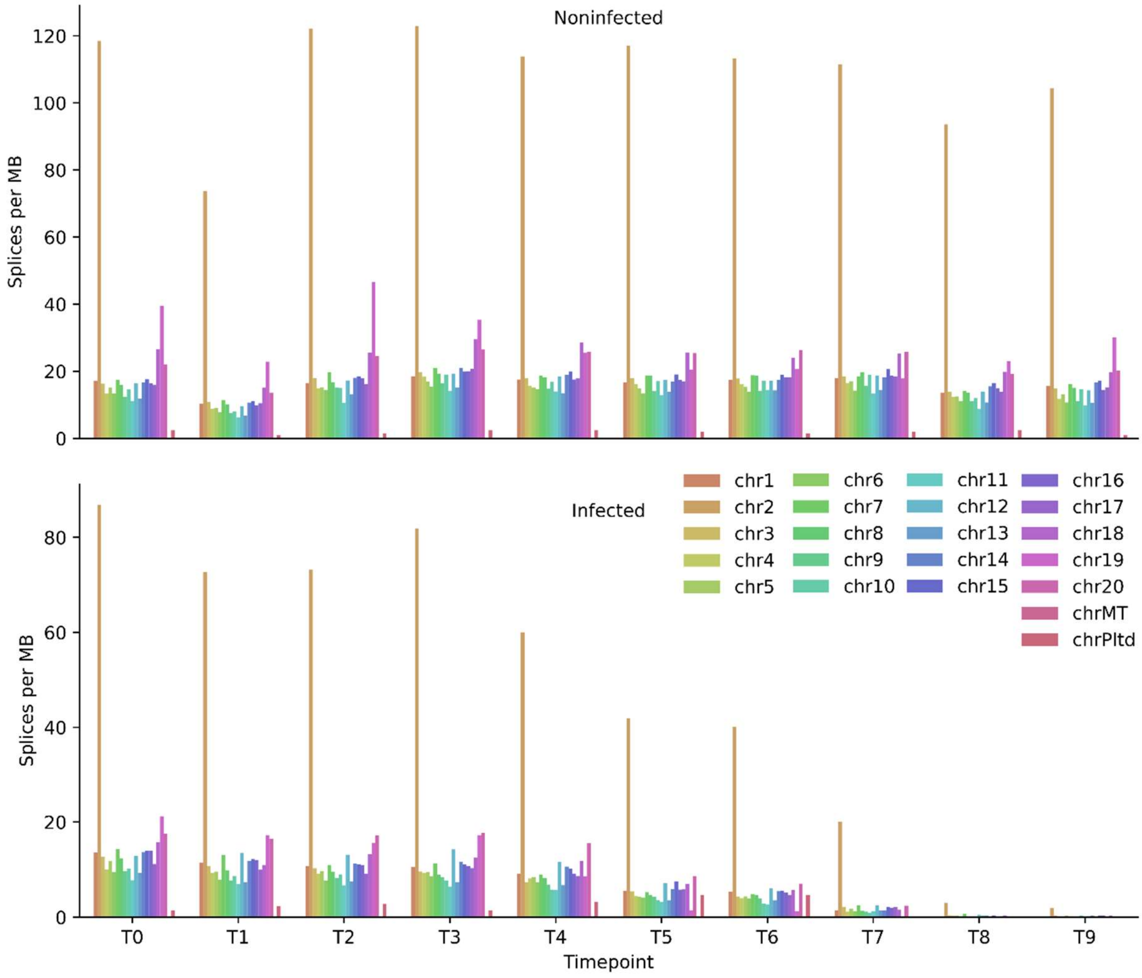


**Figure S7. Alternative Splicing across outlier regions during OtV5-infection.** (**a–b**), Splice junctions per MB across each timepoint of infection in Noninfected and OtV5-infected *O. tauri*.

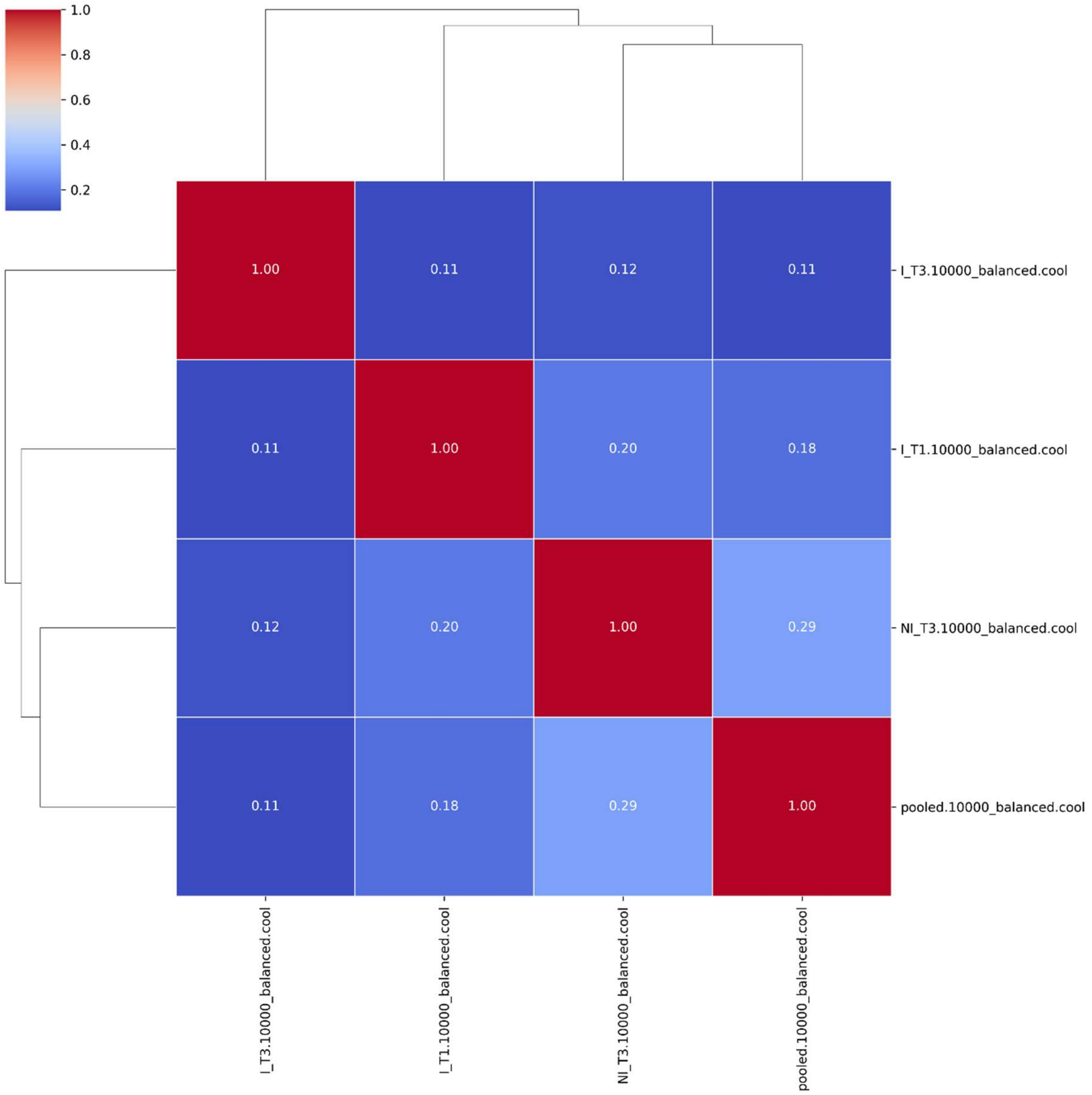


**Figure S8. Correlation between Hi-C samples.** Correlation of cool files (spearman). Note pooled is control Hi-C (Valiadi et al., 2026).

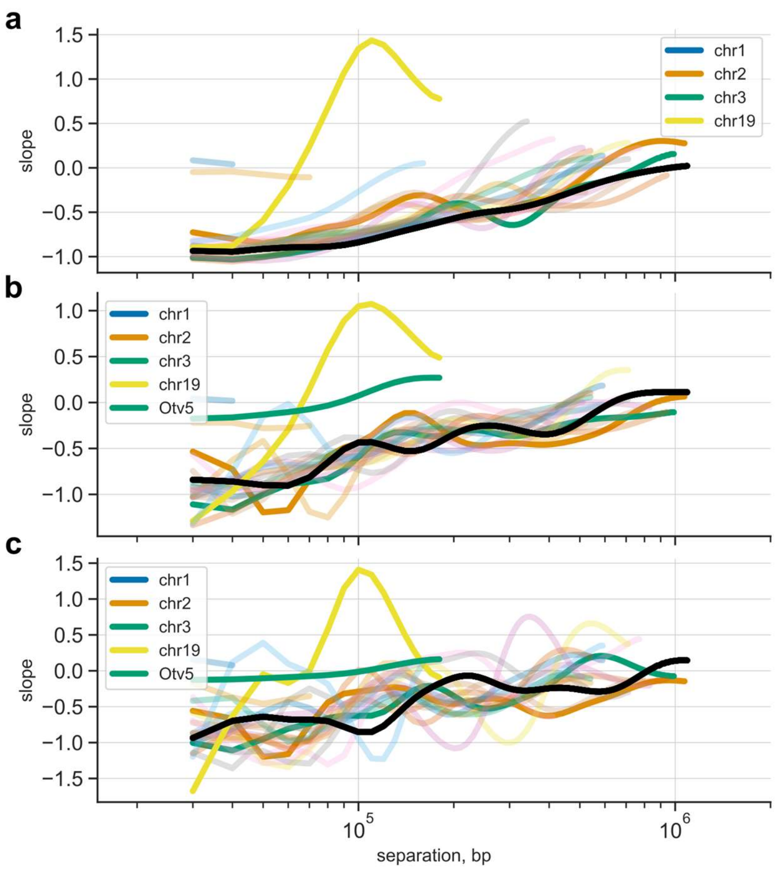


**Figure S9. Derivative of contact frequency with genomic.** (**a**), Noninfected. (**b**), 2 h p.i. (**c**), 6 h p.i.

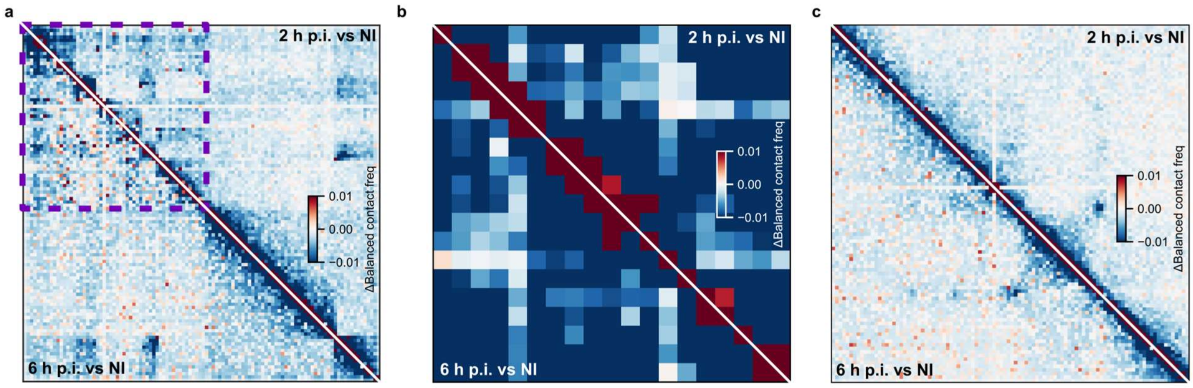


**Figure S10. Three-dimensional genome organisation of *O. tauri* during viral infection.** (**a–c**), $Log_2$ fold-change Hi-C maps showing differential contacts across infection stages: upper triangle shows 2 h p.i. vs NI, lower triangle shows 6 h p.i. vs NI, for chromosome 2 (BOC) (**a**), chromosome 19 (SOC) (**b**), and chromosome 3 (**c**).

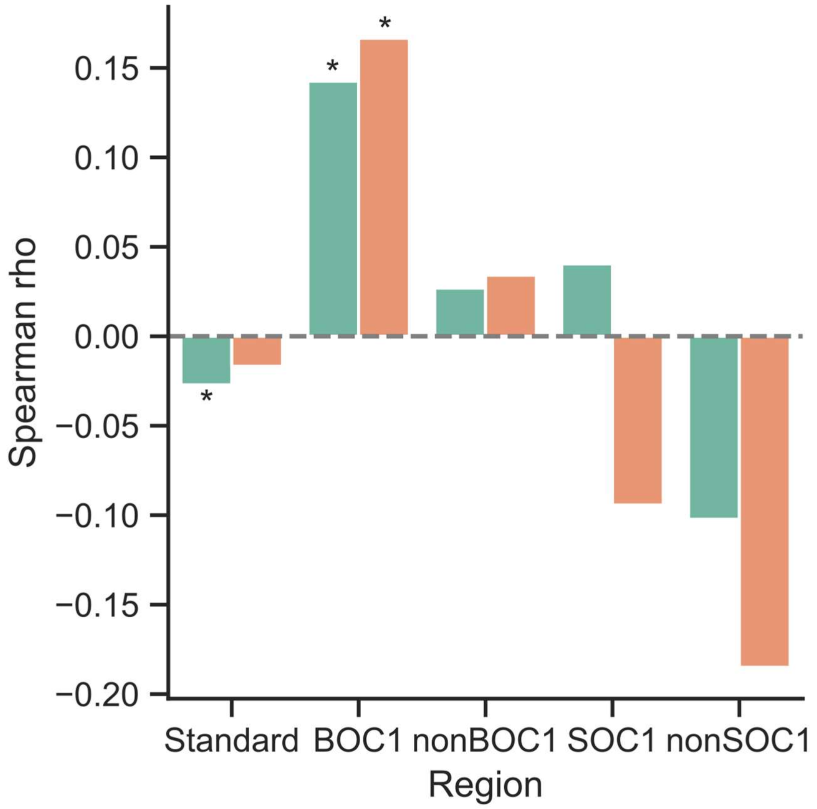


**Figure S11. BOC1 Maintains a positive correlation between gene expression and chromatin contacts throughout infection.** Spearman rank-correlation coefficients (ρ) between gene expression and host-host Hi-C contact frequency, computed across gene loci within each region at 2 hpi (green) and 6 hpi (orange). Asterisks denote significance of ρ (Spearman correlation test). Significance: *$p \leq 0.05$.

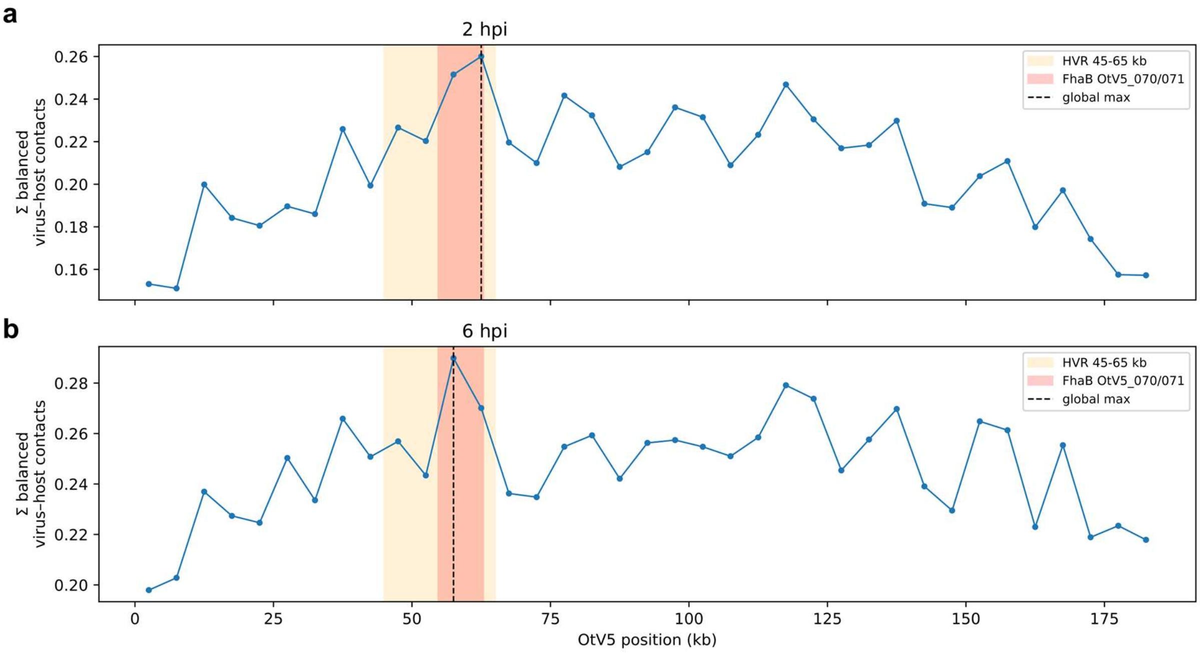


**Figure S12. Regional host–virus interaction profiles across infection timepoints.** Hi-C contact profile of OtV5 genome against host chromosomes. Balanced virus-host contacts were summed across host chromosomes in 5-kb viral bins. (**a**), T1, 2 h post-infection; (**b**), T3, 6 h post-infection, respectively. Orange shading marks the hypervariable region (HVR; 45–65 kb), and red shading marks the FhaB genes OtV5_070/OtV5_071 (54,743–62,876 bp). The dashed black vertical line indicates the globally maximal 5-kb bin in each profile.

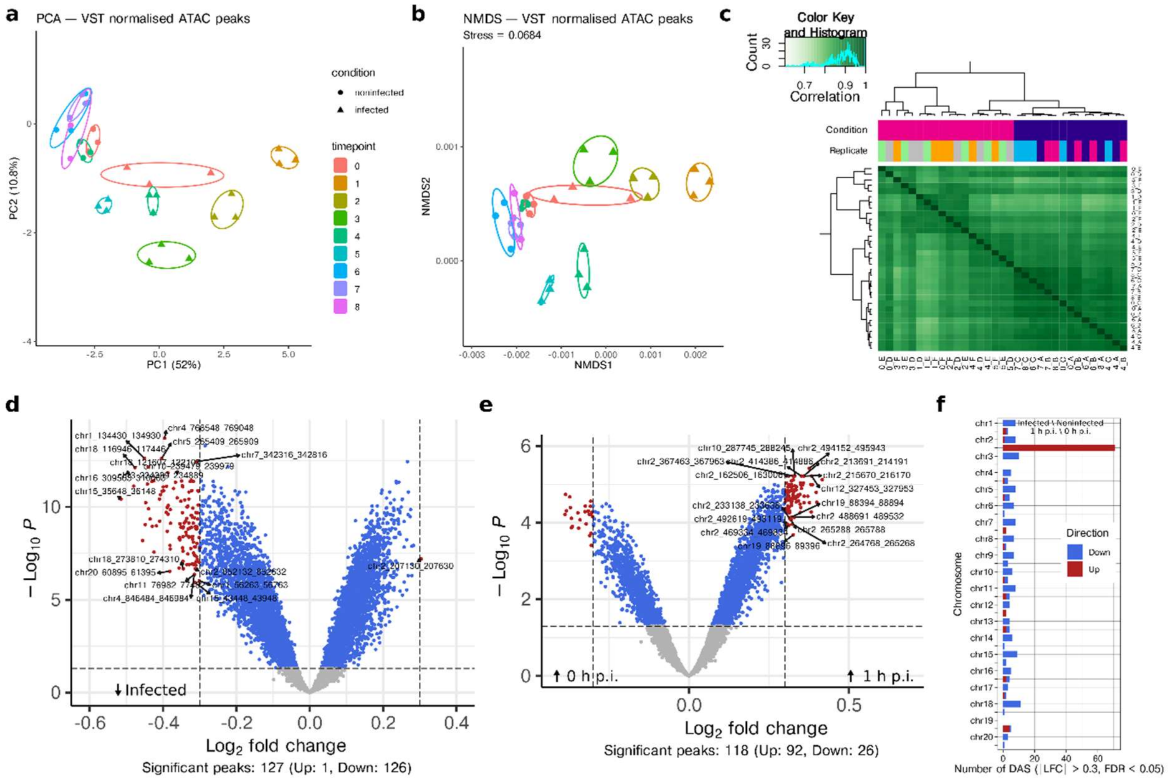


**Figure S13. ATAC-seq analysis.**

(**a**), PCA of VST-normalized ATAC peaks, with circles representing noninfected samples, filled triangles representing infected samples, and empty triangles representing noninfected replicates. (**b**), NMDS of VST-normalized ATAC peaks, same formatting as (a). (**c**), DiffBind heatmap showing clustering based on correlation between condition and replicates. (**d**), Volcano plot of infected versus susceptible cultures (whole time series design: condition + timepoint), showing differentially accessible peaks with |$Log_2$FoldChange| > 0.3 and adjusted p-value < 0.05. (**e**), Volcano plots of 1 h p.i. versus 0 h p.i. OtV5-infected cultures, showing differentially accessible peaks with |$Log_2$FoldChange| > 0.3 and adjusted p-value < 0.05. (**f**), Bar plot showing the number of differentially accessible peaks across chromosomes for infected vs noninfected and 1 h p.i. vs 0 h p.i. contrasts, with upregulated peaks in red and downregulated peaks in blue.

**Tables**

**Table S1.** Relative proportion of mean expression signal per region across noninfected transcriptomes.

| *hpi* | *BOC1* | *SOC* | *SOC1* | *Standard* |
|---|---|---|---|---|
| ***0*** | **58.83092** | **1.61216** | **24.04636** | **15.51056** |
| ***2*** | **56.36622** | **1.420436** | **27.75446** | **14.45888** |
| ***4*** | **56.00439** | **1.451002** | **26.89173** | **15.65288** |
| ***6*** | **55.6718** | **2.110773** | **19.43984** | **22.77759** |
| ***8*** | **56.6775** | **1.68484** | **19.78228** | **21.85538** |
| *10* | 58.44613 | 1.618917 | 18.91668 | 21.01827 |
| *12* | 59.16346 | 1.76795 | 17.46297 | 21.60563 |
| *14* | 58.47413 | 1.773189 | 18.28249 | 21.47019 |
| *18* | 60.27736 | 1.460018 | 26.40275 | 11.85987 |
| *24* | 57.06157 | 1.406803 | 29.33741 | 12.19422 |

**Table S2.** Relative proportion of mean expression signal per region across infected transcriptomes.

| *hpi* | *BOC1* | *SOC* | *SOC1* | *Standard* |
|---|---|---|---|---|
| ***0*** | **52.98719** | **2.321569** | **29.50095** | **15.1903** |
| ***2*** | **50.55103** | **2.169806** | **30.49148** | **16.78769** |
| ***4*** | **53.38226** | **1.711141** | **29.77774** | **15.12886** |
| ***6*** | **58.82317** | **1.418075** | **26.99367** | **12.76509** |
| ***8*** | **50.57335** | **2.796959** | **29.8694** | **16.7603** |
| *10* | 49.18272 | 3.521949 | 30.73269 | 16.56264 |
| *12* | 55.14777 | 3.906036 | 25.72876 | 15.21744 |
| *14* | 60.08725 | 3.394512 | 22.83256 | 13.68568 |
| *18* | 56.48472 | 2.116353 | 29.072 | 12.32693 |
| *24* | 47.47892 | 12.348778 | 29.89704 | 10.27526 |

**Table S3. Description of genes upregulated at 0 h and 2 h p.i. with positive SignalP classification.**

| *Gene ID* | *Description* | *0 h* | *2 h* |
|---|---|---|---|
| *OT_ostta01g04640* | Elicitor-inducible LRR receptor-like proteins | ✓ | ✓ |
| *OT_ostta01g04820* | Superoxide dismutase, copper/zinc binding domain | ✓ | ✓ |
| *OT_ostta02g05440* | 5'-Nucleotidase/apyrase | ✓ | ✓ |
| *OT_ostta03g00120* | Nucleotide-diphospho-sugar transferase | ✓ | |
| *OT_ostta03g01220* | FAS1 domain | ✓ | |
| *OT_ostta03g03140* | tRNA 4-demethylwyosine synthase | ✓ | |
| *OT_ostta03g05330* | Violaxanthin de-epoxidase (VDE) lipocalin | ✓ | |
| *OT_ostta04g00310* | Unnamed product | ✓ | |
| *OT_ostta04g01745* | Mannan endo-1,4-beta-mannosidase | ✓ | |
| *OT_ostta05g02630* | Unnamed product | ✓ | |
| *OT_ostta05g03500* | ABC-2 type transporter | ✓ | ✓ |
| *OT_ostta06g02410* | PBP domain | ✓ | |
| *OT_ostta08g01710* | Formin_Homology | ✓ | ✓ |
| *OT_ostta08g03780* | FAS1 domain | ✓ | |
| *OT_ostta10g03190* | Unnamed product | | ✓ |
| *OT_ostta11g01040* | Phospholipase/carboxylesterase/thioesterase domain-containing protein | ✓ | ✓ |
| *OT_ostta11g02250* | Peptidase S1, trypsin family, active site | ✓ | ✓ |
| *OT_ostta12g01260* | Carboxypeptidase | ✓ | |
| *OT_ostta12g01770* | Rab-GTPase-TBC domain | ✓ | |
| *OT_ostta12g02370* | Spore coat protein CotH | ✓ | |
| *OT_ostta16g02460* | alpha/beta hydrolase | ✓ | ✓ |
| *OT_ostta17g00025* | Unnamed product | ✓ | |

**Supplementary Information**

**Dataset S1 – Table of algal cell density (cells/mL) and viral-like particle abundance (VLPs/mL) in *Ostreococcus tauri* control (noninfected) and infected cultures from 0 to 24 hours post-infection.**

**Dataset S2 – Table of expression profiles (at time points 0, 2, 4, 6, and 8) and functional annotations (including EC numbers, GO terms, and FoldSeek predictions) for proteins encoded by the Ostreococcus tauri virus OtV5, with temporal classification into early, mid, and late gene clusters.**

**Dataset S3 – Table of time-course expression data (0–24 hpi) for genes of *Ostreococcus tauri* under viral infection and non-infected control conditions, including differential expression status, temporal clustering, and Gene Ontology annotations.**

**Dataset S4 – Table of enriched Gene Ontology terms for gene clusters (G1, G2, S, M) from *Ostreococcus tauri* under viral infection, including odds ratios, p-values, and FDR-adjusted significance.**

**Dataset S5 – Table of genes upregulated in *Ostreococcus tauri* at 0 hours post-infection (immediate early response) during viral infection, including log2 fold changes, significance, and protein annotations.**

**Dataset S6 – Table of genes upregulated in *Ostreococcus tauri* at 2 hours post-infection (immediate early response) during viral infection, including log2 fold changes, significance, and protein annotations.**

**Dataset S7 – Table of genes upregulated in *Ostreococcus tauri* at 6 hours post-infection (immediate early response) during viral infection, including log2 fold changes, significance, and protein annotations.**

**Dataset S8 – Table of splice junction abundance (mean, standard deviation, and mean per million per megabase) for infected and noninfected samples across timepoints (T0–T9) and chromosomes (including viral genome OtV5).**

**Dataset S9 – Table of Hi-C interaction statistics (total reads, cis/trans interactions, and short/long-range cis contacts) for *Ostreococcus tauri* under viral infection at 2 and 6 hours post-infection compared to noninfected controls.**

**Dataset S10 – Table of Spearman correlation coefficients between Hi-C interaction frequency (host-host and viral-host) at gene loci and corresponding RNA-seq expression levels during *Ostreococcus tauri* infection at 2 and 6 hours post-infection, stratified by genomic regions (SOC1, BOC1, nonBOC1, and standard).**

**Dataset S11 – Table of balanced average Hi-C contact frequencies between *Ostreococcus tauri* host chromosomes and the OtV5 viral genome at two timepoints (T1 and T3) during infection.**

**Dataset S12 – Table of viral-host Hi-C contact frequencies across 10-kb bins of the *Ostreococcus tauri* host genome at two time points (T1 and T3) during viral infection.**

**Dataset S13 – Table of the top 10 host genomic regions contacting the OtV5 virus at timepoints T1 and T3, including the viral genes (classified as early, mid, or late) associated with each region.**

**Dataset S14 – Table of viral-host Hi-C contact frequencies across 5-kb bins of the *Ostreococcus tauri* host genome at two time points (T1, 2 hpi; T3, 6 hpi), including HVR/FhaB regional mean contacts and ratios, maximal contacting bin coordinates, and permutation-based enrichment statistics.**

**Dataset S15 – Table of ATAC-seq differential accessibility peaks (Infected vs Noninfected) in *Ostreococcus tauri* during viral infection, including genomic coordinates, fold change, FDR, and associated gene annotations.**

**Dataset S16 – Table of ATAC-seq differential accessibility peaks (1 h p.i. vs 0 h p.i.) in *Ostreococcus tauri* during viral infection, including genomic coordinates, fold change, FDR, and associated gene annotations.**

**Dataset S17 – Table of Spearman correlation coefficients between ATAC-seq coverage and viral-host Hi-C contact frequency across host genomic regions (BOC1, SOC1, and standard) at 2- and 6-hours post-infection for multiple biological replicates.**
